# Neuronal substance-P drives breast cancer growth and metastasis *via* an extracellular RNA-TLR7 axis

**DOI:** 10.1101/2024.03.08.584128

**Authors:** Veena Padmanaban, Isabel Keller, Ethan S. Seltzer, Benjamin N. Ostendorf, Zachary Kerner, Sohail F. Tavazoie

## Abstract

Increased tumour innervation is associated with adverse survival outcomes in multiple cancers^1–4^. To better understand the mechanisms underlying this, we studied the impact of innervation on breast cancer metastatic progression. Metastatic mammary tumours of mice were substantially more innervated than non-metastatic isogenic tumours. Three dimensional co-cultures and *in vivo* models revealed that sensory dorsal root ganglion (DRG) neurons enhanced the growth, invasion, and systemic dissemination of cancer cells—thereby driving breast cancer metastasis. By *in vitro* screening of neuropeptides known to be secreted by DRG neurons, we identified substance-P (SP) as a mediator of these pro-metastatic functions. Neuronal SP signaled through tumoural tachykinin receptors (TACR1) to drive single-stranded RNA (ssRNA) secretion from cancer cells. Extracellular RNA acted on tumoural TLR7 receptors to activate an autocrine pro-metastatic gene expression program. In support of these findings, patient tumours with increased SP expression exhibited higher rates of lymph node metastasis. Additionally, this SP/ssRNA induced *Tlr7* gene expression signature associated with reduced breast cancer survival outcomes in two independent patient cohorts. Finally, therapeutic targeting of this neuro-cancer axis with the TACR1 antagonist aprepitant, an approved anti-nausea drug, suppressed breast cancer metastasis in multiple mouse models. Our findings reveal multiple aspects of metastatic progression to be regulated by neurons via a therapeutically targetable neuropeptide/extracellular RNA sensing axis.

## MAIN TEXT

Nerve fibers have been detected within malignant tissues for decades^5–7^. Most solid tumours receive innervation from the peripheral nervous system in the form of autonomic (sympathetic and parasympathetic) and/or sensory nerves. Perineural invasion, or the invasion of a nerve by cancer cells^8^, has been shown to correlate with poor outcome in many solid cancers^3,9–12^. An active role for tumor innervation in regulating tumourigenesis is only starting to be recognized. The autonomic nervous system has been shown to be required for tumour initiation and growth of gastric^10^ and prostate^13^ tumours. Additionally, immunomodulatory roles have recently been discovered for autonomic and sensory nerves^14,15^. We hypothesized that neural innervation may regulate metastatic progression via direct effects on cancer cells and sought to test this premise using molecular, cellular, and genetic methods.

### Sensory innervation correlates with the metastatic potential of breast tumours

To determine if expression levels of neuronal genes in human breast cancers associate with metastasis, we surveyed the expression of two pan-neuronal markers (βIII-tubulin^16^ and PGP9.5^17^) in the KM Plotter^18^ breast cancer gene expression dataset. Patients whose tumours expressed higher levels of these neuronal markers exhibited higher rates of metastatic recurrence (Extended Data Fig. 1 a-c). We next assessed the expression levels of these pan-neuronal markers in murine tumours arising from isogenic breast cancer cells of varying metastatic potential. Immunofluorescence staining and western blotting revealed that highly metastatic tumours (4T1, EO771 LM2, HCC1806 LM2) expressed higher levels of βIII-tubulin relative to poorly metastatic tumours (67NR, EO771, HCC1806 respectively; Fig. 1a, Extended Data Fig. 1d). Importantly, βIII-tubulin expression arose from the stromal compartment, consistent with its neuronal expression (Extended Data Fig. 1d). Moreover, across two independent patient cohorts totaling 30 breast tumors that we analyzed, increased innervation was associated with increased lymph node dissemination (Fig. 1 b-c). Collectively, these findings reveal that the extent of tumoural innervation correlates with metastatic propensity.

**Figure 1:**
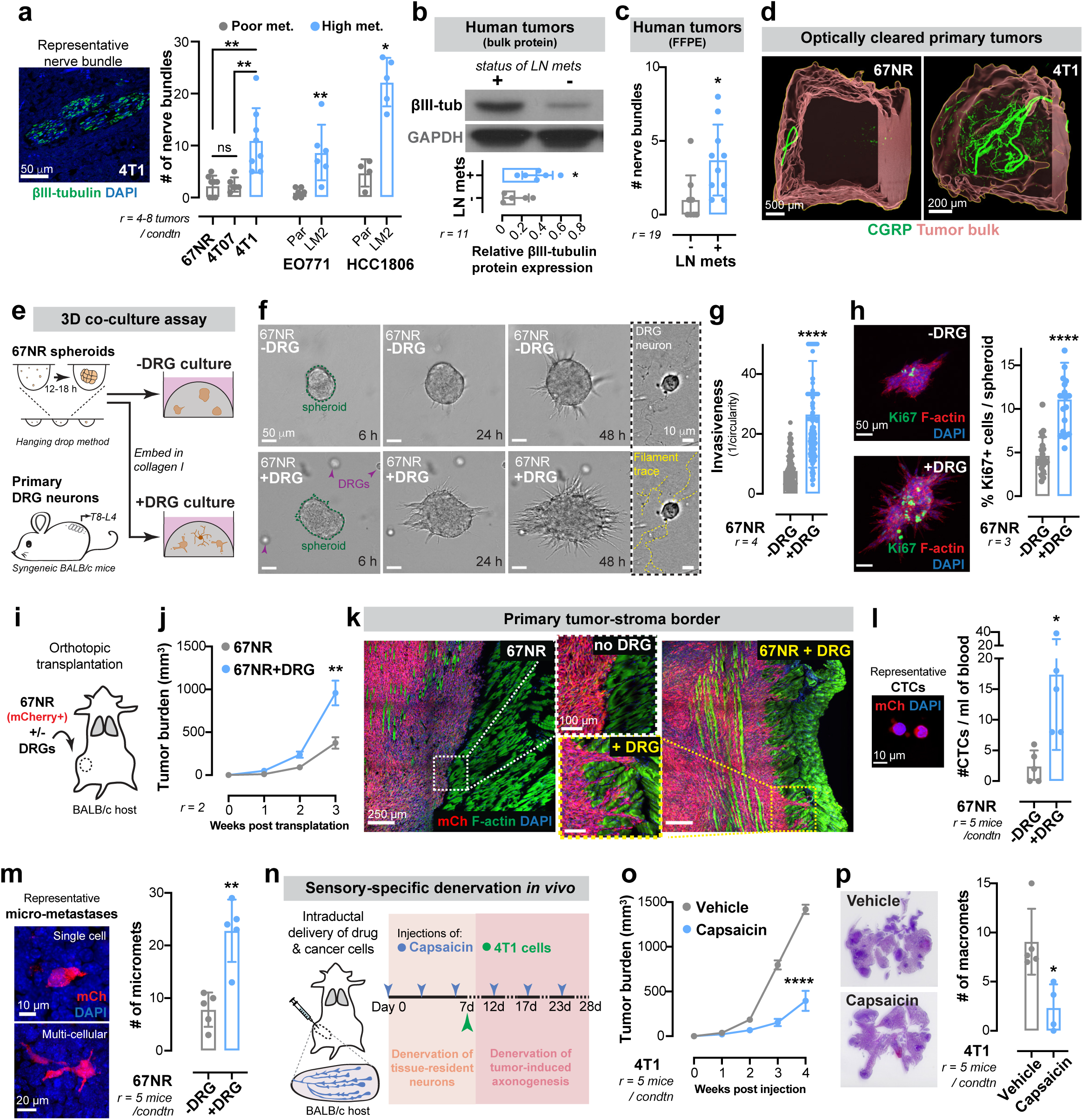
DRG neurons promote invasion, proliferation, and metastasis in breast cancer. (a) Left: Representative micrograph of a nerve bundle (marked by βIII-tubulin immunofluorescence) within a 4T1 primary tumour. Scale bar, 50 μm. Right: There is a significant increase in the number of nerve bundles in highly metatstatic primary tumours relative to corresponding isogenic poorly metastatic tumours. Data are mean ± SD. \*\**p* (67NR vs 4T1) = 0.001, \*\**p* (4T07 vs 4T1) = 0.0019, ^ns^*p* (67NR vs 4T07) = 0.9095 by one-way ANOVA; \*\**p* (EO771 Par vs LM2) = 0.0065, \**p* (HCC1806 Par vs LM2) = 0.0159 by Mann-Whitney test. (b-c) Primary human breast tumours from patients with lymph node positive disease have higher levels of βIII-tubulin compared to those with localized disease as seen in two independent cohorts by Western blotting (b; 11 patient samples; \**p* = 0.0329 by two-tailed t-test) and immunofluorescence (c; 19 patient samples; \**p* = 0.0119 by unpaired two-tailed t-test). (d) Representative micrographs showing increased CGRP+ sensory innervation in optically cleared highly metastatic 4T1 primary tumours compared to poorly metastatic 67NR primary tumours. Scale bar, 500 μm (67NR) or 200 μm (4T1). (e) Schematic of co-culture of 67NR cancer cell spheroids and primary dorsal root ganglia (DRG) neurons isolated from syngeneic BALB/c mice in 3D collagen I (invasion assay). (f) Representative micrographs of 67NR spheroids without (top row) and with (bottom row) DRG neurons. Scale bar, 50 μm. Right (dotted-line rectangle): Axons from DRG neurons branch into the collagen I matrix during the culture. Scale bar, 10 μm. (g) Co-culture with DRG neurons promotes invasiveness of 67NR spheroids. Mean ± SD. \*\*\*\**p* < 0.0001 by Mann-Whitney test. (h) Immunofluorescence showing increased proliferation of 67NR spheroids when co-cultured with DRG neurons. Scale bar, 50 μm. Mean ± SD. \*\*\*\**p* < 0.0001 by Mann-Whitney test. (i) Schematic of orthotopic transplantation of mCherry-labelled 67NR cells with or without DRG neurons into syngeneic BALB/cJ host mice. (j) Tumours arising from 67NR cancer cells co-transplanted with DRG neurons are larger than control 67NR tumours. Mean ± SEM. \*\**p* = 0.0017 by two-tailed t test. (k) Representative tile scans of primary tumours arising from 67NR cancer cells transplanted with or without DRG neurons (scale bar, 250 μm); enlarged insets of the tumour-stroma border are shown (scale bar, 100 μm). When co-injected with DRG neurons, 67NR cancer cells frequently extend collective invasion strands into the surrounding muscle. (l) Left: A representative image of mCherry+ CTCs isolated via cardiac puncture from mice bearing 67NR tumours (scale bar, 10 μm). Right: Co-transplantation with DRG neurons significantly increases the number of CTCs arising from 67NR tumours. Mean ± SD. \**p* = 0.0287 by two-sided t test. (m) Left: Representative mCherry+ lung micro-metastases in mice bearing 67NR tumours (scale bar, 10 μm). Right: Co-transplantation with DRG neurons significantly increases the number of micro-metastases from 67NR tumours. Metastases were counted in mice with size-matched tumors. Mean ± SD. \*\**p* = 0.0079 by Mann-Whitney test. (n). Schematic of sensory-specific denervation of 4T1 tumours using capsaicin. Capsaicin was injected intraductally to remove tissue-resident innervation prior to injection of 4T1 cancer cells. Capsaicin injections were continued intraductally to sustain the denervation. (o) Capsaicin-treated 4T1 tumours are significantly smaller than vehicle treated tumours. Mean ± SEM. \*\*\*\**p* < 0.0001 by two-sided t test. (p) Left: Representative H&E-stained lungs from 4T1 tumour-bearing mice treated with vehicle or capsaicin. Right: There is a significant decrease in the number of lung macro-metastases in mice treated with capsaicin. Metastases were counted in mice with size-matched tumors. \**p* = 0.0119 by two-sided t test.

To identify the source of peripheral innervation for breast tumours, we began by characterizing the pattern of innervation of the murine mammary gland. We used GFP-tagged cholera-toxin β^19^ to retrogradely label the neurons that innervate the abdominal mammary glands of mice (Extended Data Fig. 1e). We identified the dorsal root ganglia (T8-L4) sensory neurons as the predominant source of innervation for these mammary glands (Extended Data Fig. 1f). Consistently, we observed sensory innervation marked by calcitonin gene-related peptide^20^ (CGRP) within the normal ductal epithelium (Extended Data Fig. 1g). Importantly, we also observed abundant sensory innervation in syngeneic (4T1 and EO771) and genetically initiated (MMTV-PyMT and C3(1)-TAg) mouse breast cancer models (Extended Data Fig. 1h), a patient-derived model of breast cancer (Extended Data Fig. 1i), and three independent primary human breast tumours (Extended Data Fig. 1j). Further supporting sensory innervation of metastatic breast cancer, we observed a significantly higher degree of sensory innervation in optically cleared 4T1 tumours relative to isogenic poorly metastatic 67NR tumours (Fig. 1d). These findings reveal that sensory innervation is enhanced in breast tumors with higher metastatic propensity.

### Sensory neurons promote invasion, proliferation, and colony formation in breast cancer

To elucidate the mechanisms by which sensory neurons regulate tumour progression, we developed a 3D co-culture system comprised of 67NR (poorly metastatic) cancer cell spheroids and primary sensory neurons dissected from the DRGs of syngeneic naïve mice (Fig. 1e). Spheroids and neurons were embedded in 3D collagen I, a protein that is highly enriched in the stroma of breast cancer^21^. During the culture, DRG neurons extended axons into the surrounding matrix (Fig. 1f). When cultured in the presence of DRG neurons, 67NR breast cancer spheroids exhibited significantly more invasion (Fig. 1 f-g) and proliferation (Fig. 1h). Consistent with these *in vitro* findings, cancer cells adjacent to nerve bundles in 4T1 primary tumours were more proliferative than cancer cells from a distant region of the same tumour (Extended Data Fig. 2a). In a Matrigel-based colony formation assay, DRG neurons enhanced the colony forming capacity of 67NR breast cancer cells in a cell-dose dependent manner (Extended Data Fig. 2 b,c). DRGs similarly increased the invasiveness and colony formation capacity of MMTV-PyMT derived organoids and cancer cell clusters, respectively (Extended Data Fig. 3 a-c). We next investigated the effect of DRGs on primary human tumour organoids by isolating tumour organoids from four distinct patients’ breast tumours (3/4 ER+ PR+ HER2- and 1/4 ER+ PR+ HER2+) and co-cultured them with murine DRG neurons (Extended Data Fig. 3d). Consistent with our findings in mouse models, DRG neurons strongly increased invasion and proliferation of organoids from all four human tumours (Extended Data Fig. 3 e-f). These findings reveal that DRG neurons enhance the proliferative and invasive phenotypes of murine and human breast cancer cells *in vitro*.

We next assessed the impact and requirement of *in vivo* sensory innervation on breast cancer metastasis. To do this, we sought to increase innervation of poorly metastatic 67NR tumours and conversely to denervate highly metastatic 4T1 tumours. We first co-transplanted a small number of DRG neurons (∼200) along with mCherry+ 67NR cancer cells into syngeneic BALB/cJ hosts (Fig. 1i). Tumoral mCherry expression allowed us to track cancer cells within non-fluorescent hosts as they progressed through the metastasis cascade. We observed a significant increase in the extent of sensory innervation in the co-transplantation condition (Extended Data Fig. 4a), suggesting that DRG neurons are functional when injected *in vivo*. Co-transplantation with sensory neurons significantly increased the growth of 67NR primary tumours (Fig. 1j). Careful examination of the tumour-stroma border revealed that DRGs promoted the collective invasion of 67NR cancer cells into the surrounding muscle (Fig. 1k). It is widely believed that 67NR cancer cells fail to metastasize due to their inability to intravasate^22^. However, the use of a fluorescent reporter and imaging at single-cell resolution revealed the presence of hundreds of 67NR cancer cells within the lungs of mice (Extended Data Fig. 4 b-d). A vast majority of these cells were associated with the lung endothelium, consistent with a failure of these cells to metastasize due to impaired extravasation (Extended Data Fig. 4 b,c). A minority of 67NR cells had exited the lung endothelium to form micro-metastases (Extended Data Fig. 4d). This contrasted with isogeneic 4T07 cells, which efficiently formed micro-metastases, and 4T1 cells, which efficiently formed macro-metastases^22^. Mice co-transplanted with 67NR cells and DRGs harbored a significantly higher number of circulating mCherry positive tumour cells (Fig. 1l) and micro-metastases (Fig. 1m). Additionally, in an experimental metastasis assay, co-injection of 67NR cancer cells with DRG neurons significantly increased metastatic colonization (Extended Data Fig. 4e-g). Consistent with this metastasis-promoting role for sensory neurons, we also observed enhanced tumour growth and micro-metastases in co-transplantation experiments performed with EO771 breast cancer cells (Extended Data Fig. 3 g-i). These findings reveal that DRG neurons are sufficient to enhance the metastatic capacity of breast cancer cells in co-transplantation experiments *in vivo*.

In a complementary set of experiments, we performed sensory-specific denervation of 4T1 tumours using capsaicin, a TRPV1 agonist and neurotoxin that degenerates primary sensory nerves^23^ (Extended Data Fig. 5a). We first injected capsaicin into the mammary ductal tree to remove tissue-resident sensory innervation, then intraductally injected 4T1 cancer cells while continuing to administer capsaicin to maintain a sustained denervated state (Fig. 1n). By performing capsaicin injections intraductally, we restricted its effects within the ductal tree. We discovered that sensory-specific denervation greatly reduced tumour growth (Fig. 1o) and suppressed the number of macro-metastases (quantified in mice bearing size-matched primary tumours; Fig. 1p). Importantly, capsaicin had no effect on the invasiveness or proliferation of 4T1 cancer cells *in vitro*, consistent with the impact of capsaicin on these phenotypes occurring via its effects on neurons, and not on the tumour compartment itself (Extended Data Fig. 5 b-d). To further control for possible confounding effects of TRPV1 agonism in immune cells^24^, we repeated these experiments in NOD-SCID gamma mice bearing 4T1 tumours and observed similar anti-tumour effects of capsaicin (Extended Data Fig. 5 e-f). Taken together, these results reveal a requirement for sensory innervation for optimal breast cancer progression and metastasis.

### Neuronal substance-P drives breast cancer metastasis

A significant route for metastasis is the migration of cancer cells along nerves, a process called perineural invasion^8^. This, however, requires some physical contact between neurons and cancer cells. In our co-culture models, despite strong tumourigenic effects of DRG neurons, physical interaction between these cell types were not evident. We therefore hypothesized that DRG neurons may mediate pro-metastatic effects on cancer cells via secreted molecules. To directly test this, we harvested conditioned media from DRG-67NR co-cultures (DRG-CM; Fig. 2a). We used conditioned media from 67NR only cultures as control (tumour CM). Treatment with DRG-CM phenocopied the enhanced invasive and proliferative effects of DRG neurons (Figure 2 b,c). These results suggest that a secreted factor(s) mediates the pro-tumourigenic effects of DRG neurons.

**Figure 2:**
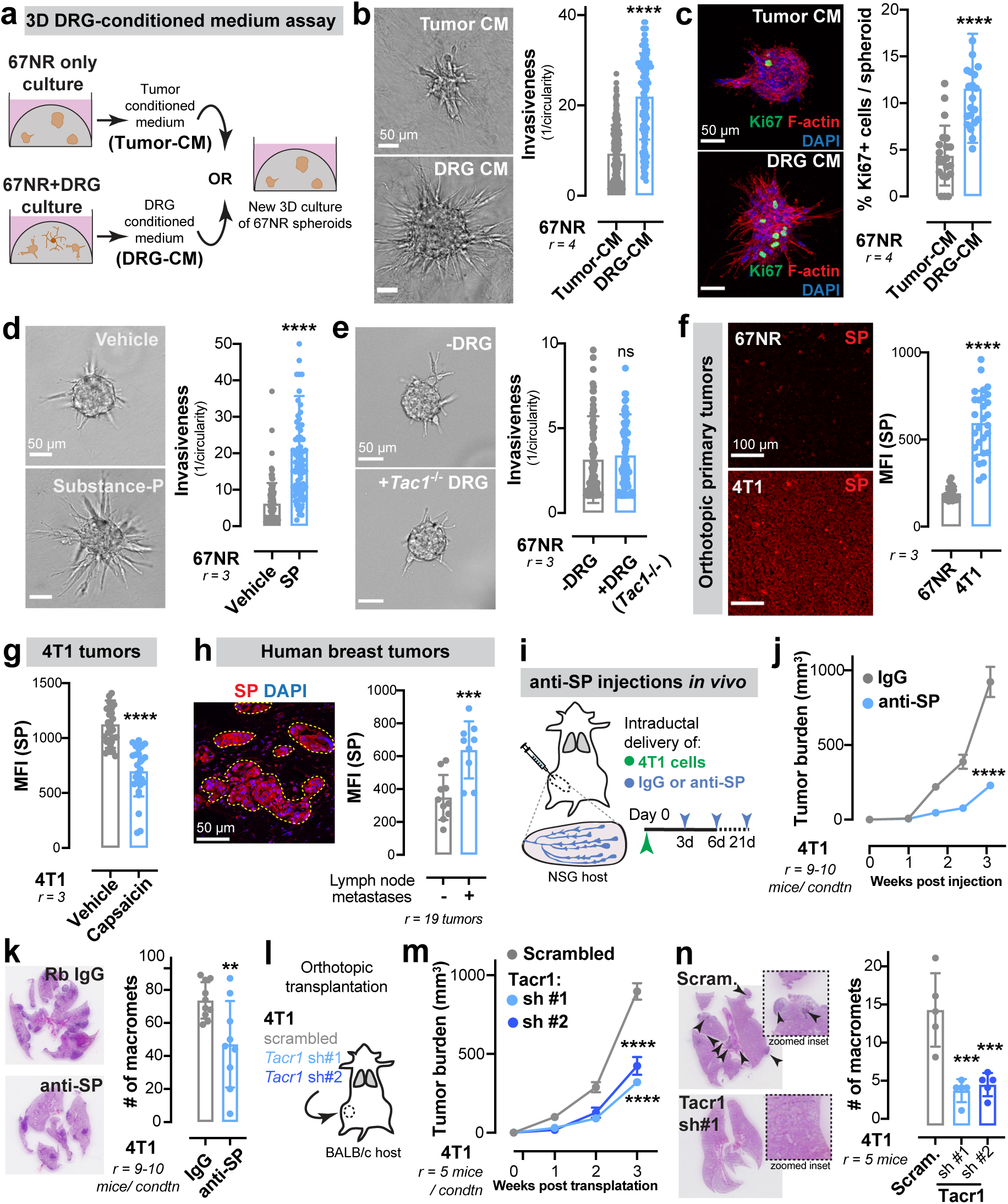
DRG-derived substance-P promotes breast cancer metastasis. (a) Schematic for isolation of conditioned medium. Tumour-CM is isolated from a 3D culture of 67NR spheroids in collagen I. DRG-CM is isolated from a 3D co-culture of 67NR spheroids and DRG neurons. (b) DRG-CM significantly promotes the invasiveness of 67NR spheroids. Scale bar, 50 μm. Mean ± SD. \*\*\*\**p* < 0.0001 by Mann-Whitney test. (c) Immunofluorescence showing increased proliferation of 67NR spheroids cultured with DRG-CM. Scale bar, 50 μm. Mean ± SD. \*\*\*\**p* < 0.0001 by Mann-Whitney test. (d) Treatment with substance-P (SP) significantly increases the invasiveness of 67NR spheroids. Scale bar, 50 μm. Mean ± SD. \*\*\*\**p* < 0.0001 by Mann-Whitney test. (e) DRG neurons isolated from *Tac1*-null mice do not promote invasiveness of 67NR spheroids. Scale bar, 50 μm. Mean ± SD. ^ns^*p* = 0.3070 by Mann-Whitney test. (f) Left: Immunofluorescence showing increased SP in 4T1 primary tumours compared to 67NR tumours. Scale bar, 50 μm. Right: Mean fluorescence intensity (MFI) of SP ± SD. \*\*\*\**p* < 0.0001 by unpaired two-tailed t-test. (g) Capsaicin treated 4T1 tumours express significantly lower SP compared to vehicle treated tumours. Mean ± SD. \*\*\*\**p* < 0.0001 by unpaired two-tailed t-test. (h) Left: Representative micrograph of a primary human breast tumour stained for SP with the tumour region marked as a ROI used for MFI calculations. Right: SP expression is significantly higher in breast tumours of patients with lymphatic spread compared to those with localized disease. Mean ± SD. \*\*\**p* = 0.0008 by unpaired two-tailed t-test. (i) Following intraductal injections of 4T1 cancer cells, serial injections of IgG control or SP blocking antibody were performed in NSG mice. (j) Treatment with an anti-SP antibody significantly decreases the size of 4T1 tumours. Mean ± SEM. \*\*\*\**p* < 0.0001 by Mann-Whitney test. (k) Left: Representative H&E-stained lungs from 4T1 tumour-bearing mice treated with IgG or an anti-SP antibody. Right: There is a significant decrease in the number of lung macro-metastases in mice treated with an anti-SP antibody. Metastases were counted in mice with size-matched tumors. Mean ± SD. \*\**p* = 0.0092 by two-sided t test. (l) 4T1 cancer cells transduced with either scrambled control, or two independent shRNAs targeting *Tacr1* were orthotopically transplanted into the abdominal mammary glands of syngeneic BALB/cJ hosts. (m) Depletion of TACR1 significantly decreased the growth of 4T1 tumours. Mean ± SEM. \*\*\*\**p* < 0.0001 by one-way ANOVA. (n) Left: Representative H&E-stained lungs from 4T1 tumour-bearing mice transduced with scrambled or Tac1r shRNA hairpins. Arrows: metastases. Right: There is a significant decrease in the number of lung macro-metastases in mice treated with an anti-SP antibody. Metastases were counted in mice with size-matched tumors. Mean ± SD. \*\*\**p* = 0.0003 by one-way ANOVA.

Sensory DRG neurons secrete neuropeptides that stimulate various biological processes. We tested the ability of three common DRG-secreted neuropeptides, CGRP^25^, substance-P (SP)^26^, and galanin^27^ to promote invasion of 67NR spheroids into collagen I. We found that SP and galanin significantly increased 67NR spheroid invasion, with SP exerting the largest magnitude effect, while CGRP had a modest effect (Fig. 2d, Extended Data Fig. 6a). Importantly, DRG-CM had significantly higher levels of SP compared to tumour-CM (Extended Data Fig. 6b). As an orthogonal approach, we performed a co-culture containing DRG neurons that do not synthesize SP (isolated from T*ac1 null* mice^28^, which are deficient for the T*ac1* gene that encodes the pre-peptide that is processed to generate SP) and 67NR cancer cells. We found that T*ac1*^-/-^ DRG neurons were unable to impact the invasiveness of 67NR spheroids (Fig. 2e), revealing neuropeptide SP to be required for the pro-invasive effects of DRG neurons on breast cancer cells.

To assess the impact of neuronal-SP on metastasis *in vivo*, we first quantified SP levels in 67NR and 4T1 primary tumours. We found that mice bearing highly metastatic 4T1 tumours had higher SP expression in the tumour (Fig. 2f) and in plasma (Extended Data Fig. 6c), relative to mice with poorly metastatic 67NR tumours. This SP expression was innervation-dependent since intraductal treatment of 4T1 tumours with capsaicin, a sensory neurotoxin, significantly reduced tumoural SP levels (Fig. 2g). Importantly, in a cohort of 19 primary human breast tumours, SP expression was higher in tumours of patients with lymph node metastases relative to those with localized disease (Fig. 2h). To directly test the requirement for SP in breast cancer growth and metastasis, we intraductally administered an anti-SP blocking antibody to 4T1 tumour-bearing mice (Fig. 2i, Extended Data Fig. 6d). Inhibition of extracellular SP strongly reduced tumour growth and metastasis (quantified in mice bearing size-matched primary tumours; Fig. 2 j,k). SP mediates neurotransmission by acting on the tachykinin receptor 1 (TACR1 or NK1R)^29^. To determine if SP mediates pro-metastatic phenotypes via TACR1, we used lentiviral shRNA-mediated silencing to deplete Tacr1 in 4T1 cancer cells (Extended Data Fig. 6e). Tacr1 depletion significantly reduced 4T1 spheroid invasion and proliferation *in vitro* (Extended Data Fig. 6 f,g) and 4T1 tumour growth and metastasis *in vivo* (metastases were quantified in mice bearing size-matched primary tumours; Fig. 2 l-n). Taken together, these data identify neuronal substance-P as a driver of breast cancer progression and metastasis.

### Substance-P promotes the release of pro-metastatic ssRNA molecules from cancer cells

Surprisingly, we discovered that conditioned medium from DRG neurons alone (DRG-only-CM) contained less SP than conditioned medium from the DRG-67NR co-culture described above (DRG-CM, Extended Data Fig. 6b) and was not sufficient to increase the invasiveness of 67NR spheroids (Extended Data Fig. 6 h-j). These data suggest that in addition to neuronal SP, one or more additional tumoural factor(s) are present in DRG-CM that are (i) produced upon the interaction of sensory neurons with cancer cells and (ii) required to promote breast cancer invasiveness. To identify such factor(s), we treated DRG-CM with DNase, RNase A, or heat inactivated its protein components. While DNase treatment and heat inactivation had little effect on the potency of DRG-CM (Extended Data Fig. 7a), RNase A treatment significantly impaired the ability of DRG-CM to promote invasion of 67NR spheroids (Fig. 3 a-b). Using RNases specific for single-stranded RNA (ssRNA, RNase T1^30^) or double stranded RNA (dsRNA, RNase III^31^), we implicated ssRNAs as mediators of the pro-invasive effects of DRG-CM on breast cancer cells (Fig. 3 a-b).

**Figure 3:**
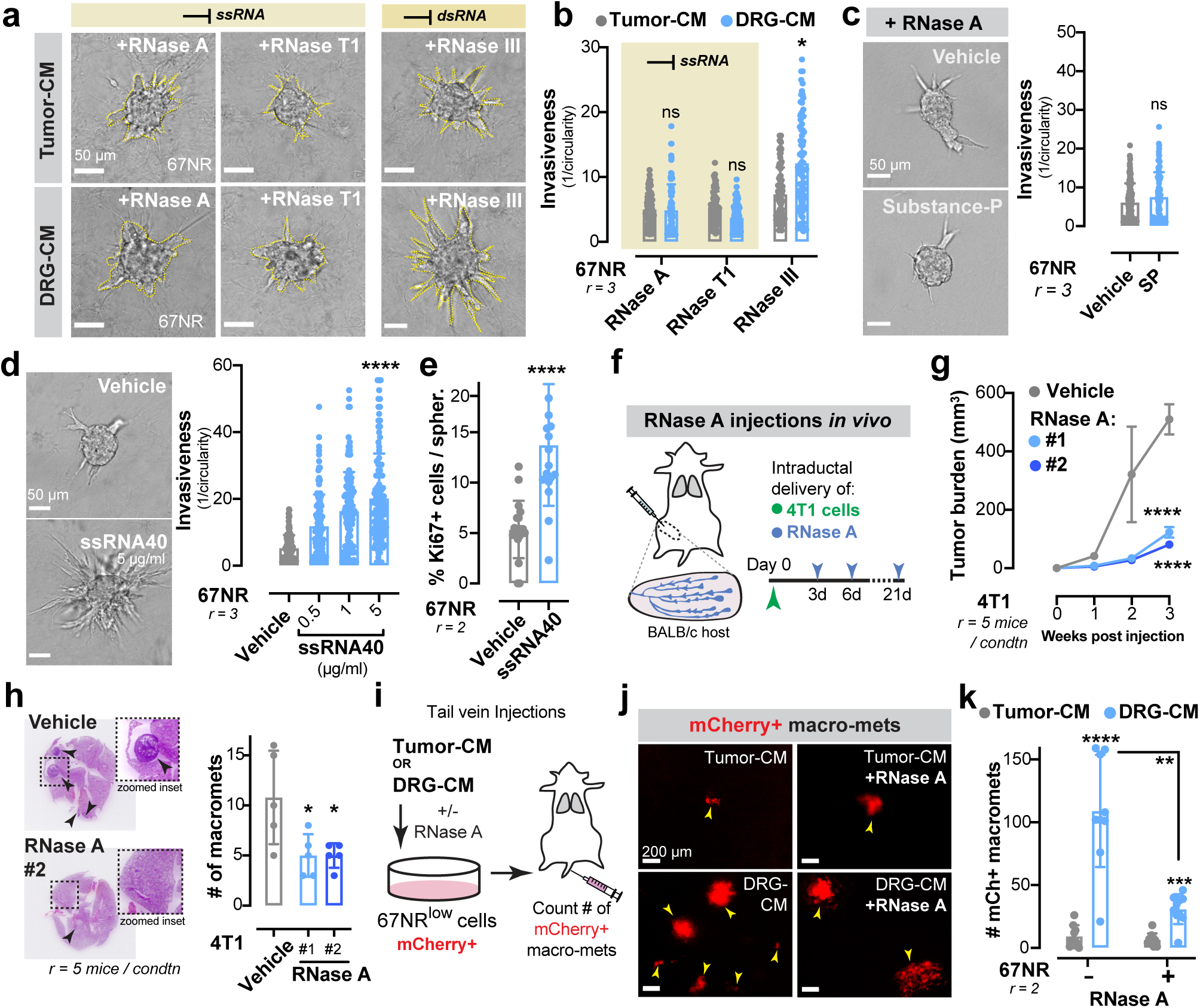
DRG-derived SP drives the secretion of pro-metastatic ssRNA molecules. (a) Representative micrographs of 67NR spheroids cultured in tumour-CM (top row) or DRG-CM (bottom row) in the presence of either RNase A, RNase T1, or RNase III. Scale bar, 50 μm. (b) Treatment with RNase A and RNase T1 (ssRNA specific degradation) significantly inhibits the potential of DRG-CM to drive invasiveness of 67NR spheroids. Mean ± SD. ^ns^*p* > 0.9999, \**p* = 0.0215 by Kruskal-Wallis test. (c) SP treatment in the presence of RNase A has no effect on the invasiveness of 67NR spheroids. Scale bar, 50 μm. Mean ± SD. ^ns^*p* = 0.2416 by Mann-Whitney test. (d-e) Addition of a ssRNA mimetic to the culture medium significantly increases 67NR spheroid invasion (d) and proliferation (e) in a dose-dependent manner. Scale bar, 50 μm. Mean ± SD. \*\*\*\**p* < 0.0001 by Kruskal-Wallis test (d) and Mann-Whitney test (e). (f) 4T1 cancer cells were injected intraductally, followed by injections with vehicle or two independent RNase A enzymes every 3 days for 3 weeks. (g) RNase A treatment of 4T1 tumours significantly decreases tumour size. Mean ± SEM. \*\*\*\**p* < 0.0001 by one-way ANOVA. (h) Left: Representative H&E-stained lungs from 4T1 tumour-bearing mice treated with vehicle or RNase A. Arrows: metastases. An enlarged region is shown in the inset. Right: There is a significant decrease in the number of lung macro-metastases in mice treated with RNase A. Metastases were counted in mice with size-matched tumors. Mean ± SD. \**p* = 0.0213 by one-way ANOVA. (i) mCherry+ 67NR cancer cells were pre-treated with tumour-CM or DRG-CM with or without RNase A for 48 hours prior to tail vein injections into syngeneic BALB/cJ host. (j) Representative micrographs of mCherry+ (67NR) lung metastases counted 1-week post-injection. Arrows: metastases. (k) Pre-treatment of 67NR cancer cells with DRG-CM increased the number of mCherry+ metastases compared to those pre-treated with tumour-CM. Addition of RNase A to DRG-CM significantly decreased its metastasis-promoting ability. Mean ± SD. \*\*\*\**p* < 0.0001, \*\*\**p* = 0.0005, \*\**p* = 0.0015 by Mann-Whitney test.

Thus far, we have shown that both SP and ssRNAs in DRG-CM are required for promoting cancer spheroid invasion and proliferation. We next sought to determine if SP and ssRNAs signal through a common pathway that drives breast cancer metastasis. We observed that treatment of 67NR spheroids with SP in the presence of RNase A completely abolished the invasion-promoting effects of SP (Fig. 3c). Since this experiment was performed using exogenously added SP and in the absence of sensory neurons, we propose a model wherein sensory neurons secrete SP, which in turn promotes the release of ssRNA molecules from breast cancer cells that drive invasion and proliferation.

To further define the role of ssRNA in breast cancer metastasis, we treated 67NR spheroids with a ssRNA mimetic, ssRNA40^32^. Treatment with ssRNA40 significantly increased spheroid invasion and proliferation (Fig. 3 d-e). In contrast, the dsRNA mimetic Poly I:C^33^ did not confer these phenotypes (Extended Data Fig. 7b). Moreover, intraductal injections of two independently generated recombinant RNase A enzymes significantly impaired tumour growth and metastasis of 4T1 cancer cells without any observed impact on dsRNA levels (Fig. 3 f-h, Extended Data Fig. 7c).

We have thus far shown that DRG-conditioned medium exerts pro-tumourigenic effects *in vitro*. To evaluate the impact of DRG-conditioned medium on metastasis *in vivo*, we pre-treated 67NR cancer cells with DRG-CM or tumour-CM for two days prior to tail vein injection into syngeneic mice (Fig. 3i). Pre-treatment with DRG-CM was sufficient to confer metastasis-forming capacity onto the poorly metastatic 67NR cells, demonstrated by the formation of hundreds of macro-metastases (Fig. 3 j-k). Importantly, pre-treatment of 67NR cancer cells with DRG-CM containing RNase inhibited the ability of DRG-CM to confer a metastasis promoting phenotype (Fig. 3 j-k). Collectively, these data uncover a signaling axis whereby sensory nerves secrete SP, which in turn promotes release of ssRNA species from breast cancer cells that drive metastasis. In further support of this model, we found that SP and ssRNA40 promoted the invasiveness of spheroids from two additional murine breast cancer cell lines (4T1 and PyMT-derived Py8119 cells) and the human breast cancer cell line (MDA-MB-231; Extended Data Fig. 8 a,b).

### Sensory neuron-mediated signaling in cancer cells occurs via Tlr7

Our findings thus far reveal that extracellular ssRNA species can enhance the invasiveness of breast cancer cells and promote breast cancer metastasis. The detection of ssRNAs is mediated by the Tlr7 receptor in mice, which performs innate immune RNA sensing and effector responses^34^. To determine if tumoural Tlr7 mediates the downstream pro-metastatic effects of ssRNAs, we used RNA-interference to deplete Tlr7 in 67NR cancer cells and then co-cultured them with primary DRG neurons (Fig. 4 a,b; Extended Data Fig. 9a). DRG neurons were unable to promote invasiveness or proliferation of 67NR spheroids depleted of Tlr7 (Fig. 4 c-f). In contrast, 67NR spheroids lacking the dsRNA sensor Tlr3^33^ co-cultured with DRGs retained their invasiveness (Extended Data Fig. 9 b-d). Additionally, lentiviral shRNA-mediated depletion of Tlr7 in 4T1 cancer cells reduced spheroid invasion *in vitro* (Fig. 4g, Extended Data Fig. 9e), and tumour growth and metastasis *in vivo* (Fig. 4 h-j). Importantly, RNase A treatment of tumours arising from Tlr7-depleted 4T1 cells did not further reduce tumour growth (Extended Data Fig. 9 g-i), consistent with a common pathway comprising ssRNA/Tlr7. Taken together, these data reveal that neuron-dependent release of ssRNA species from cancer cells signals in an autocrine/paracrine manner via Tlr7 to drive invasion, growth, and metastasis.

**Figure 4:**
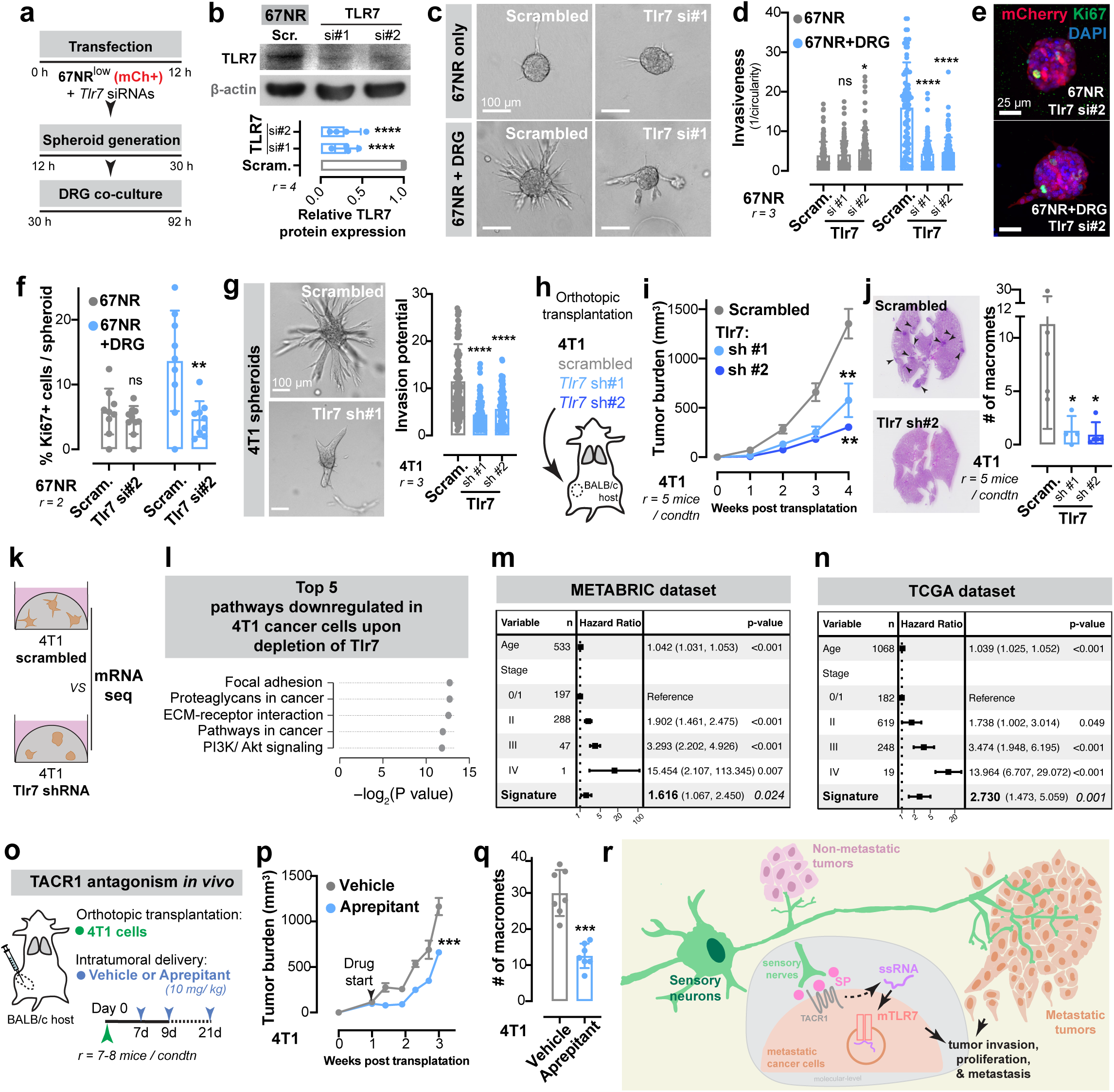
DRG neurons promote metastasis via TLR7 signaling in breast cancer cells. (a) Workflow for the transfection of 67NR cancer cells with siRNAs targeting Tlr7, generation of spheroids, and co-culture with DRG neurons. (b) Successful knockdown of TLR7 upon siRNA transfection in 67NR cancer cells was validated by Western blotting. Mean ± SD. \*\*\*\**p* < 0.0001 by one-way ANOVA. (c) Representative micrographs of 67NR spheroids (scrambled and *Tlr7*-targeting siRNA) cultured with or without DRG neurons. Scale bar, 100 μm. (d) TLR7 depletion in 67NR cancer cells does not affect its invasiveness at baseline. DRG neurons lose their potential to promote 67NR invasion upon TLR7 depletion. Mean ± SD. ^ns^*p* = 0.5787, \**p* = 0.031, ****p < 0.0001 by one-way ANOVA. (e) Representative micrographs of 67NR spheroids transfected with scrambled or siRNA targeting *Tlr7*, co-cultured with DRG neurons, and immuno-stained with Ki67. Scale bar, 25 μm. (f) Loss of TLR7 expression in 67NR spheroids significantly decreases proliferation of 67NR spheroids cultured with DRG neurons. Mean ± SD. ^ns^*p* = 0.8751, \*\**p* = 0.0048 by unpaired two-tailed t-test. (g) Left: Representative micrographs of 4T1 spheroids transduced with either scrambled or Tlr7 shRNA (two independent hairpins) in 3D collagen I. Scale bar, 100 μm. Right: Loss of TLR7 significantly decreases the invasion of 4T1 spheroids. \*\*\*\**p* < 0.0001 by Kruskal-Wallis test. (h) 4T1 cancer cells transduced with scrambled or *Tlr7*-targeting shRNAs were orthotopically transplanted into the mammary fat pad of BALB/cJ hosts. (i) Loss of TLR7 expression significantly decreased the size of 4T1 tumours. Mean ± SEM. \*\**p* = 0.0070 (scrambled vs sh#1), 0017 (scrambled vs sh#2) by one-way ANOVA. (j) Loss of TLR7 expression significantly decreased the number of macro-metastases arising from 4T1 tumours. Metastases were counted in mice with size-matched tumors. \*\**p* = 0.00390 by one-way ANOVA. (k) mRNA sequencing analysis was performed by comparing the transcriptomes of control and *Tlr7* depleted 4T1 breast cancer spheroids embedded in 3D collagen I. (l) Top five downregulated pathways in 4T1 spheroids depleted for *Tlr7* as assessed by gene set enrichment analysis (*P* values according to permutation testing). (m-n) Multivariate analysis of the association of age, tumour stage and a *Tlr7*-dependent gene signature with survival in breast cancer patients of the METABRIC (m) and TCGA (n) studies (*P* values according to multivariate Cox proportional hazard models, error bars indicate 95% confidence intervals; *n* = 533 and 1068 for METABRIC and TCGA datasets, respectively). (o) Aprepitant, a clinically used anti-memetic, was evaluated for its potential in inhibiting breast cancer progression and metastasis. 4T1 cancer cells were orthotopically transplanted into the mammary fat pad of syngeneic BALB/cJ mice. Intratumoural aprepitant injections were performed every ∼2-3 days for ∼ 3 weeks. (p) Aprepitant treatment significantly decreases 4T1 tumour growth. Mean ± SEM. \*\*\**p* = 0.0003 by Mann-Whitney test. (q) Aprepitant treatment significantly decreases the number of metastases arising from 4T1 tumours. Metastases were counted in mice with size-matched tumors. Mean ± SD. \*\*\**p* = 0.0006 by Mann-Whitney test. (r) Model figure: Breast tumours are frequently innervated by sensory nerves. Metastatic breast tumours have more sensory innervation than isogeneic tumours with lower metastatic potential. In the tumour stroma, sensory nerves secrete substance-P (SP) that activates TACR1 signaling in cancer cells that results in the secretion of ssRNA molecules. These ssRNA molecules activate TLR7 signaling on cancer cells in an autocrine/ paracrine manner to drive breast cancer metastasis.

Tlr7 is the predominant sensor for ssRNAs derived from pathogens ^35^. To define the tumoural gene expression response downstream of Tlr7 activation, we performed mRNA sequencing of control and Tlr7-depleted 4T1 spheroids (Fig. 4k). Gene-set enrichment analysis (GSEA) revealed that Tlr7 depletion repressed expression of genes implicated in cancer cell invasion, proliferation, and metastasis—most notably focal adhesions, ECM-receptors, and PI3K-Akt signaling pathway components (Fig. 4l). We calculated a gene expression signature based on differential gene expression upon Tlr7 depletion and used it to stratify breast cancer patient survival. In multivariate analyses conducted across both the TCGA and the METABRIC^36^ cohorts, we found that the Tlr7-high signature correlated with poorer overall survival (TCGA: HR 2.730, *p* = 0.001; METABRIC: HR 1.616, *p* = 0.024; Fig. 4 m,n). These findings reveal that Tlr7 promotes a pro-invasive and proliferative gene expression program in breast cancer cells that associates with poor breast cancer patient survival.

### Perturbation of sensory neuron-cancer signaling with an antiemetic inhibits metastasis

Our results implicate neuronal SP and its receptor TACR1 in driving breast cancer metastatic progression. Aprepitant is a small molecule therapeutic drug that acts as an antagonist of TACR1^37^ and is used clinically to treat nausea^38^. *In vitro*, aprepitant significantly decreased invasiveness of 4T1, Py8119, and MDA-MB-231 spheroids in a dose-dependent manner (Extended Data Fig. 10 a). Administration of clinically relevant doses of aprepitant to mice significantly inhibited tumour growth and metastatic progression in multiple models including 4T1 cancer cells (Fig. 4 o-q), Py8119 cancer cells (Extended Data Fig. 10 b,c), and MMTV-PyMT organoids (Extended Data Fig. 10 b,d). Taken together, our findings identify a neuro-cancer axis comprising SP/TACR1/ssRNA/Tlr7 as a key driver of breast cancer progression and metastasis. We also provide proof-of-concept for therapeutic targeting of multiple nodes in this axis, including an approved anti-nausea medication. Given the safety and tolerability of aprepitant, which is now available in generic form for the treatment of nausea, our findings warrant investigations into the clinical efficacy of this agent upon its prolonged use in breast cancer in combination with standard of care regimens for the prevention of metastatic relapse.

Nerves are beginning to be recognized as critical signaling structures within the tumour microenvironment. Recent studies have revealed tumourigenic^10,13,15,39,40^ and tumour suppressive^15,41,42^ functions of the autonomic nervous system. The mammary gland receives abundant sensory fiber input^43,44^, yet the impact of this sensory innervation on breast cancer progression is poorly characterized. Conceptually, our work implicates sensory neurons as drivers of tumour invasion, growth, and metastasis in breast cancer. We found that neuronal substance-P elicits the release of ssRNAs from cancer cells, which act in an autocrine manner on tumoural TLR7 to promote invasion and metastasis (Fig. 4o)—thus linking neural signaling to RNA release and an innate RNA sensing response in the tumour compartment. Furthermore, we discovered an instructive role for cancer cells in altering neuronal signaling to support cancer progression. Similar reprogramming of neurons was recently observed in head and neck cancer^45^. Taken together, we describe an elaborate signaling crosstalk between sensory neurons and cancer cells that reinforces the emerging concept of neuronal contribution to cancer progression.

Neurons are evolutionarily conserved modulators of immunity and their immune-regulatory functions have been shown to be critical in tumorigenesis^14,15,46,47^. In contrast to these previously reported indirect effects on the tumour stroma, we observed pro-tumourigenic effects of innervation directly on cancer cells via the activation of tumoural TLR7 signaling. While TLR7 canonically functions in innate-immunity as a ssRNA pathogen sensor^35^, we uncovered an immune-independent, tumour-intrinsic, and metastasis-associated transcriptional response upon neuronally-induced TLR7 activation in these cancer cells. The ability of cancer cells to selectively exploit tumourigenic responses downstream of innate immune receptors while repressing immune activating responses has been previously described^48,49^. Our findings provide proof-of-concept for targeting this sensory neuro-cancer circuit as a promising anti-metastatic approach and motivate further testing of the anti-mimetic aprepitant in human breast cancer.

## ACKOWLEDGEMENTS

We are grateful to members of our laboratory for critical discussions and feedback on the manuscript text. We thank the various resource centers at Rockefeller University: A. North, C. Pyrgaki and staff of the Bio Imaging Resource Center, C. Zhao and staff of the Genomics Resource Center, and S. Mazel and staff of the Flow Cytometry Resource Center. The results published here are in part based upon data generated by the TCGA Research Network. This work was supported by U54CA261701, R35CA274446, the Black Family Metastasis Center, the Breast Cancer Research Foundation and the Reem Kayden award. V.P. was supported by the Hope Funds for Cancer Research postdoctoral fellowship. B.N.O. was supported by a Max Eder grant of the German Cancer Aid (reference 70114327) and is a fellow of the digital clinician scientist program at BIH-Charité.

## AUTHOR CONTRIBUTIONS

V.P. and S.F.T. conceptualized the study, designed experiments, supervised research, and wrote the manuscript with input from all authors. V.P. performed most experiments with assistance from I.K., Z.K., and M.K. B.N.O. analyzed mRNA sequencing data. S.F.T. obtained funding and supervised scientists.

---

**Extended Data Figure 1:**
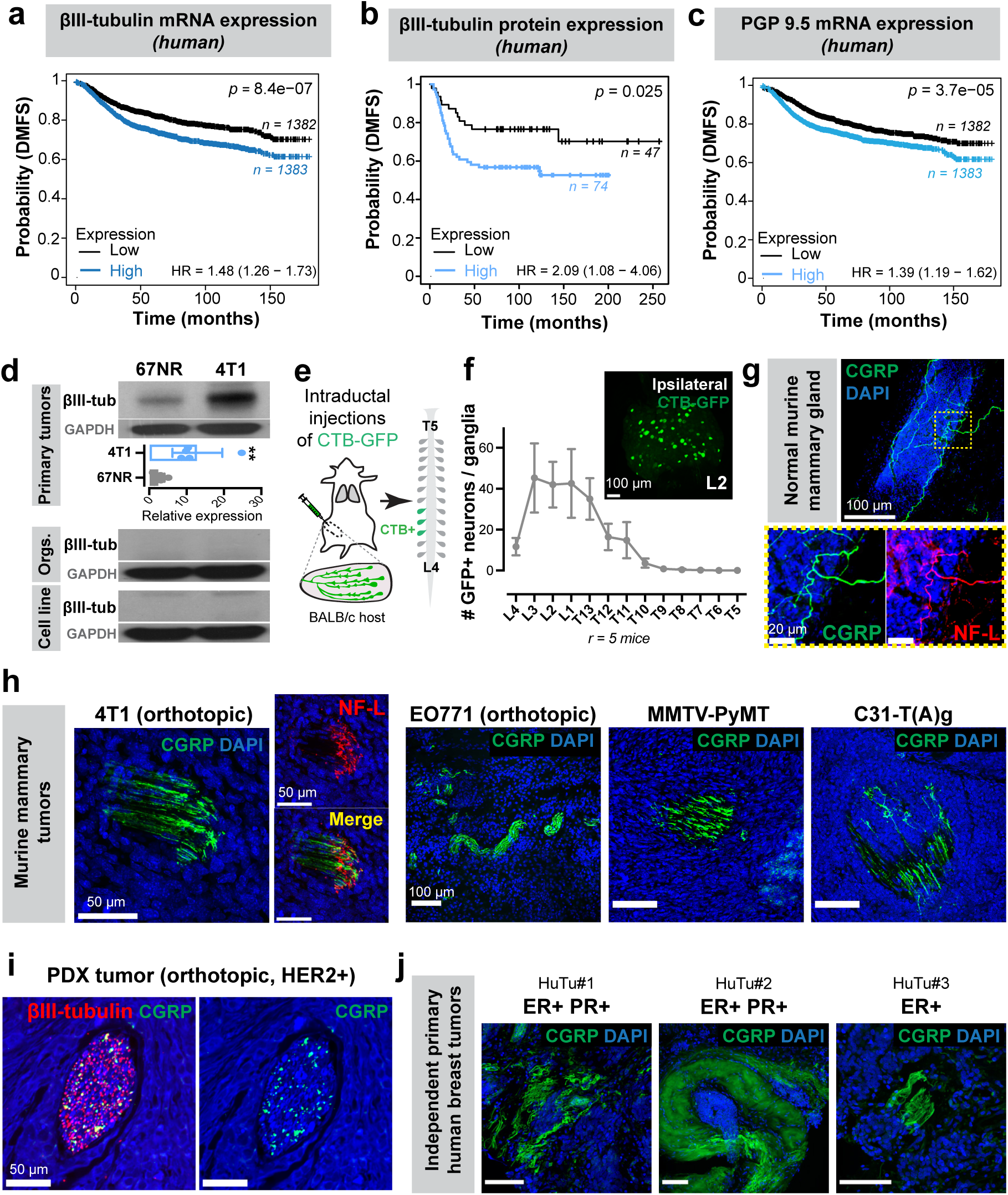
Breast tumours are frequently innervated by sensory nerves. (a-c) Kaplan-Meier plots showing distant metastasis-free survival (DMFS) of breast cancer patients sorted by the median expression level of βIII-tubulin (mRNA (a), protein (b)) or PGP9.5 (mRNA (c)) of their tumours. The x-axes in (a) and (c) were terminated at 180 months to have at least ∼10 surviving patients in each arm. (d) Representative Western blot showing higher βIII-tubulin expression in highly metastatic 4T1 tumours relative to poorly metastatic 67NR tumours. Importantly, no βIII-tubulin expression was detected in these cell lines in culture or from epithelial tumour organoids isolated from the respective primary tumours, consistent with the expression of βIII-tubulin within innervating nerves. Mean ± SD. \*\**p* = 0.0048 by unpaired two-tailed t-test. (e) GFP-tagged cholera toxin β (CTB-GFP) was intraductally injected into BALB/cJ mice to retrograde label neurons innervating the murine abdominal mammary glands. (f) Neurons within the dorsal root ganglion (DRG), largely from T12-L4 ganglia consist of neurons that innervate the abdominal mammary glands of mice. Top right: Representative L2 ganglion from the ipsilateral side of a female BALB/cJ mouse injected with CTB-GFP. Scale bar, 100 μm. (g-j) CGRP+ sensory innervation is present in normal mammary glands (g), murine mammary tumours including – orthotopic (4T1, EO771 cell-line derived tumours), and spontaneous models (MMTV-PyMT and C3(1)-TAg) (h), a patient-derived xenograft (PDX) tumour (i), and across three independent primary human tumours (j). Scale bar, 50 μm or 100 μm as noted in the corresponding figure panels.

**Extended Data Figure 2:**
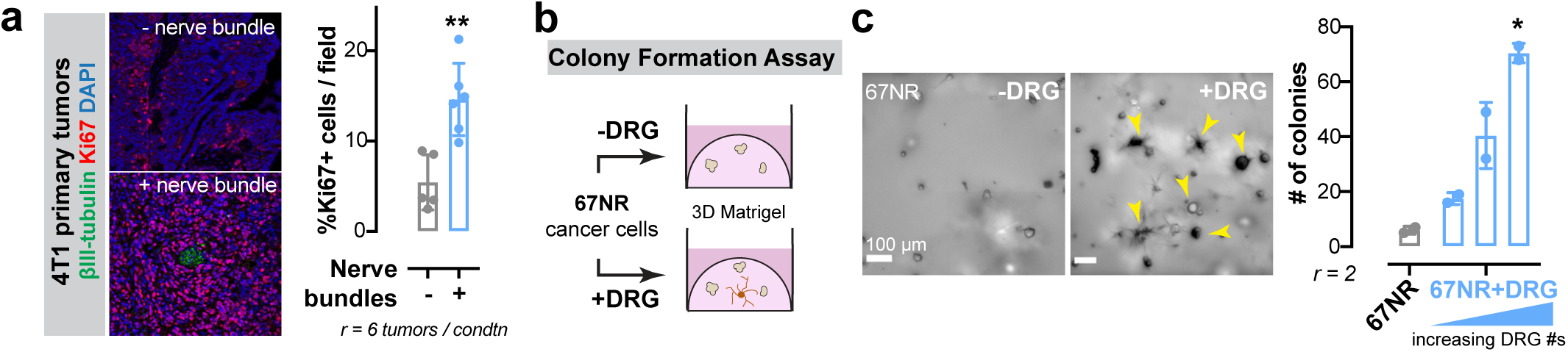
DRG neurons promote proliferation and colony formation in breast cancer. (a) Left: Representative micrographs of 4T1 primary tumours showing mitotically active cells (Ki67+). The top panel represents a tumour region with no nearby nerve bundles. The bottom panel represents a distinct region within the same tumour with a nerve bundle. Right: There is a significant increase in the percentage of Ki67+ cancer cells when adjacent to a nerve bundle. Mean ± SD. \*\**p* = 0.0043 by Mann-Whitney test. (b) Schematic for a 3D colony formation assay. 67NR cancer cells were cultured alone or in the presence of DRG neurons in 3D Matrigel. The number of colonies (>7 cells) were counted after 4 days in culture. (c) Left: Representative micrographs of the colony formation assay using 67NR cells with or without DRG neurons. Arrows: colonies. Scale bar, 100 μm. Right: Presence of DRG neurons increases colony formation by 67NR cancer cells in a cell-dose-dependent manner. Mean ± SD. \**p* = 0.0429 by Kruskal-Wallis test.

**Extended Data Figure 3:**
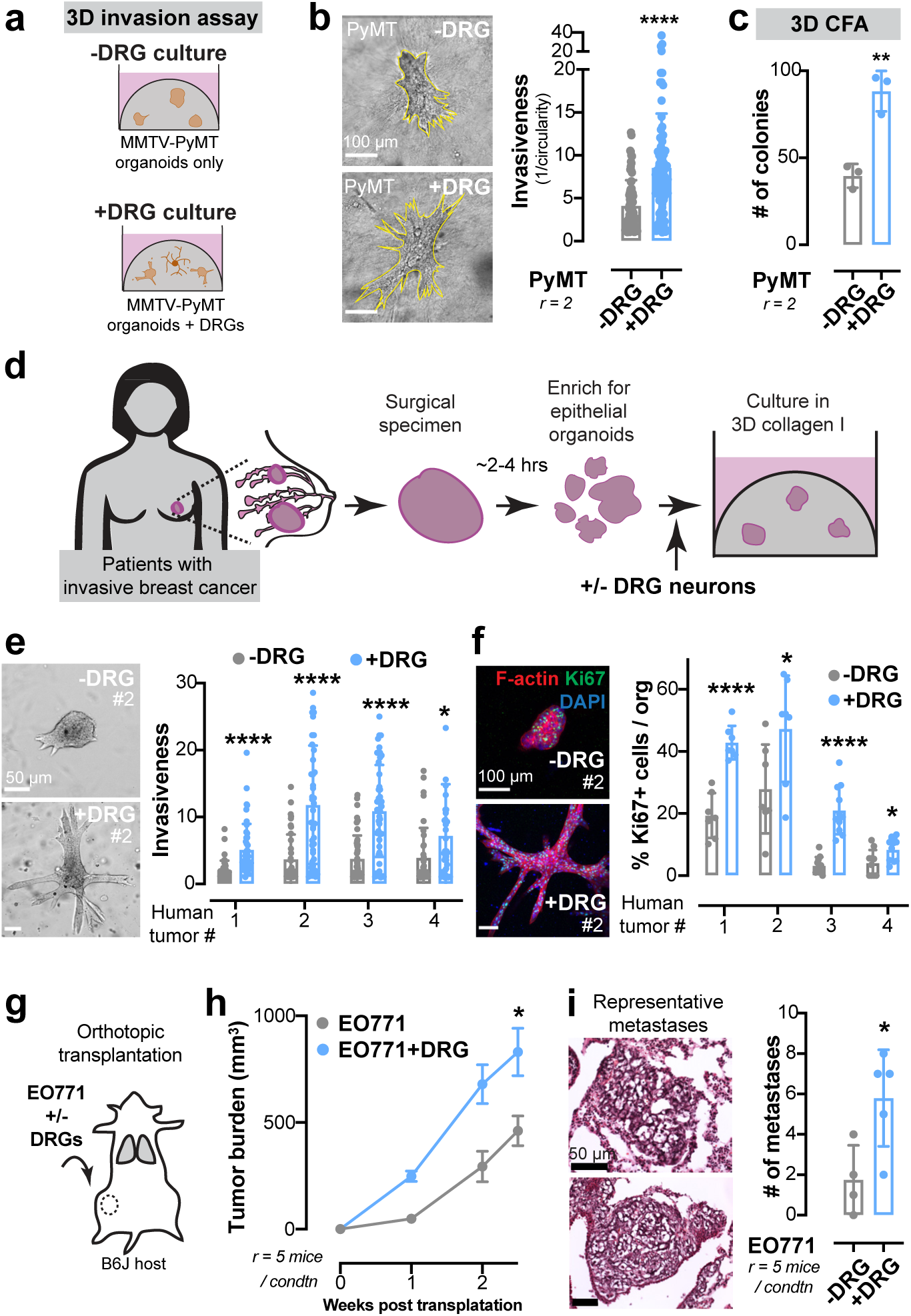
DRG neurons promote invasion, proliferation, and metastasis across multiple murine and human models of breast cancer. (a) Organoids, or clusters of ∼100-300 cancer cells, were isolated from late-stage MMTV-PyMT tumours. They were embedded either alone or with DRG neurons in 3D collagen I. (b) Left: Representative micrographs of invasion assay using PyMT organoids with or without DRG neurons. Right: DRG neurons promote invasiveness of MMTV-PyMT organoids. Mean ± SD. \*\*\*\**p* < 0.0001 by Mann-Whitney test. (c) Co-culture with DRG neurons significantly increases colony formation from MMTV-PyMT cancer cell clusters. Mean ± SD. \*\**p* = 0.0033 by unpaired two-tailed t-test. (d) Schematic for the isolation of organoids from surgical samples of primary human breast tumours. Surgical samples were processed physically and enzymatically to generate epithelial tumour organoids that were embedded in 3D collagen I. For co-cultures, human tumour organoids were cultured alongside with primary DRG neurons isolated from female C57B6/J mice. (e) Left: Representative micrographs of human tumour organoids isolated from the same primary tumour and cultured without (top) or with (bottom) DRG neurons. Right: DRG neurons significantly increase the invasiveness of primary human tumour organoids, as seen across all the 4 tumour samples. Mean ± SD. \*\*\*\**p* < 0.0001, \**p* = 0.0266 by Mann-Whitney test. (f) Left: Representative confocal images of human tumour organoids isolated from the same primary tumour and cultured without (top) or with (bottom) DRG neurons. Right: DRG neurons significantly increase the proliferation of primary human tumour organoids, as seen across all the 4 tumour samples. Mean ± SD. \*\*\*\**p* < 0.0001, \**p* = 0.0401 (#2), 0.0267 (#4) by unpaired two-tailed t-test. (g) EO771 cancer cells were orthotopically injected with or without DRG neurons into the abdominal mammary gland of syngeneic C57B6/J mice. (h) EO771 cancer cells co-injected with DRG neurons form significantly larger tumours compared to those injected without DRGs. Mean ± SEM. \**p* = 0.0345 by unpaired two-tailed t-test. (i) Left: Representative metastases positively identified in the lungs of tumour bearing mice using H&E staining. Right: Co-transplantation with DRG neurons significantly promotes formation of metastases from EO771 tumours. Mean ± SD. \**p* = 0.0249 by unpaired two-tailed t-test.

**Extended Data Figure 4:**
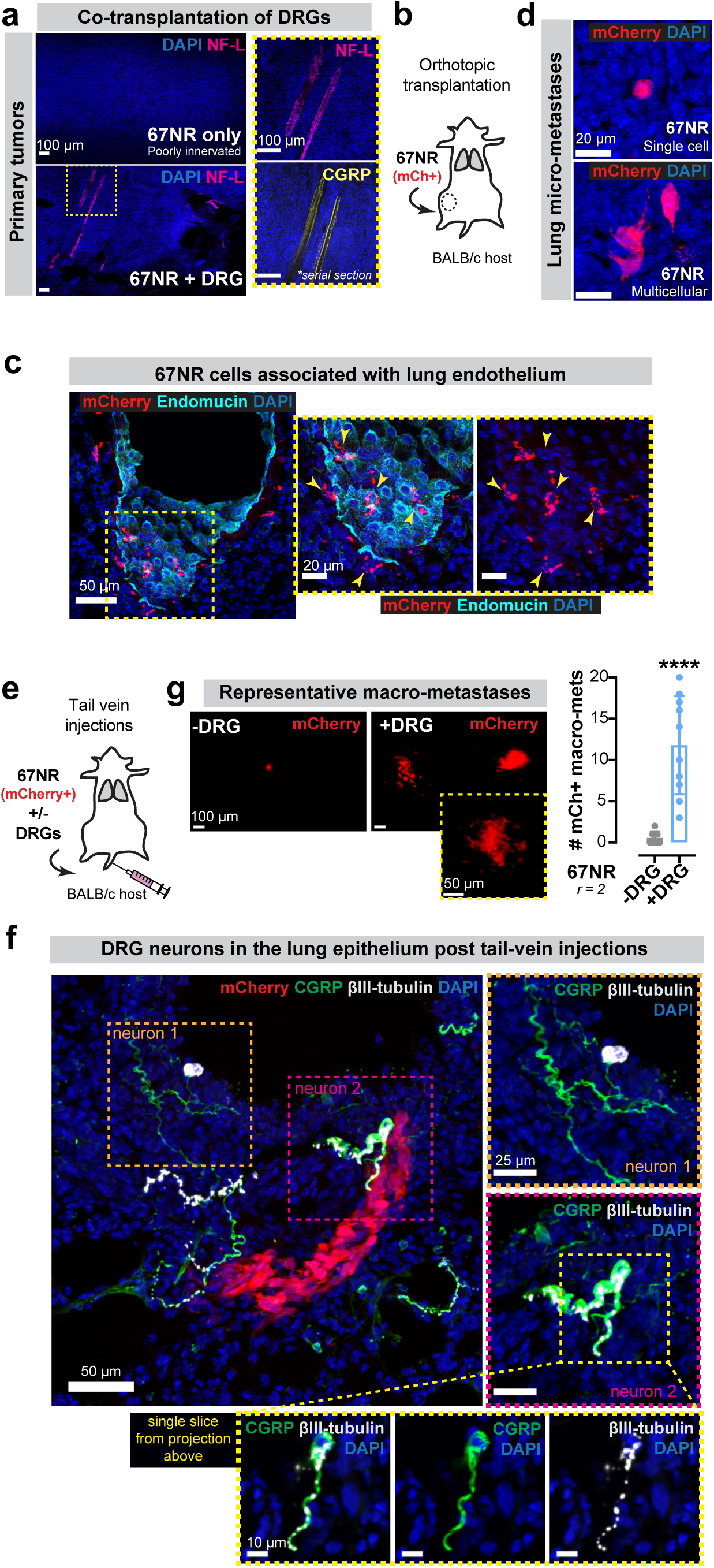
Co-injection with DRG neurons promotes survival and metastatic colonization of 67NR cancer cells. (a) Tile-scan of representative regions within 67NR primary tumours when transplanted alone (top) or with DRG neurons (bottom). Co-transplanted tumours have significantly more sensory innervation (CGRP+), consistent with the injection of DRG neurons. Scale bar, 100 μm. (b) mCherry-labeled 67NR cancer cells were orthotopically injected into syngeneic BALB/cJ host mice. (c) Following transplantation into the mammary gland, mCherry+ 67NR cancer cells are frequently associated with the lung endothelium (marked by endomucin). Arrows: mCherry+ cells. Scale bar, 50 μm. (d) Representative micrometastases in the lungs of mice transplanted with mCherry+ 67NR cancer cells. Scale bar, 20 μm. (e) Schematic of tail vein injections of mCherry-labelled 67NR cancer cells with or without primary DRG neurons into syngeneic BALB/cJ mice. (f) Representative micrographs from lungs of mice co-injected with 67NR cancer cells and DRG neurons. Lung metastases are mCherry+. Two independent neuronal cell bodies (each zoomed into individual insets to the right; scale bar, 25 μm.) can be positively identified using pan-neuronal (βIII-tubulin) and sensory (CGRP) nerve markers, consistent with the survival of these neurons when injected *in vivo* via the tail vein. Scale bar, 50 μm. (g) Left: Representative micrographs of mCherry+ lung metastases counted 1-week post-injection. Right: Co-injection of DRG neurons with 67NR cancer cells via the tail vein of mice significantly increased metastatic lung colonization. Mean ± SD. \*\*\*\**p* < 0.0001 by Mann-Whitney test.

**Extended Data Figure 5:**
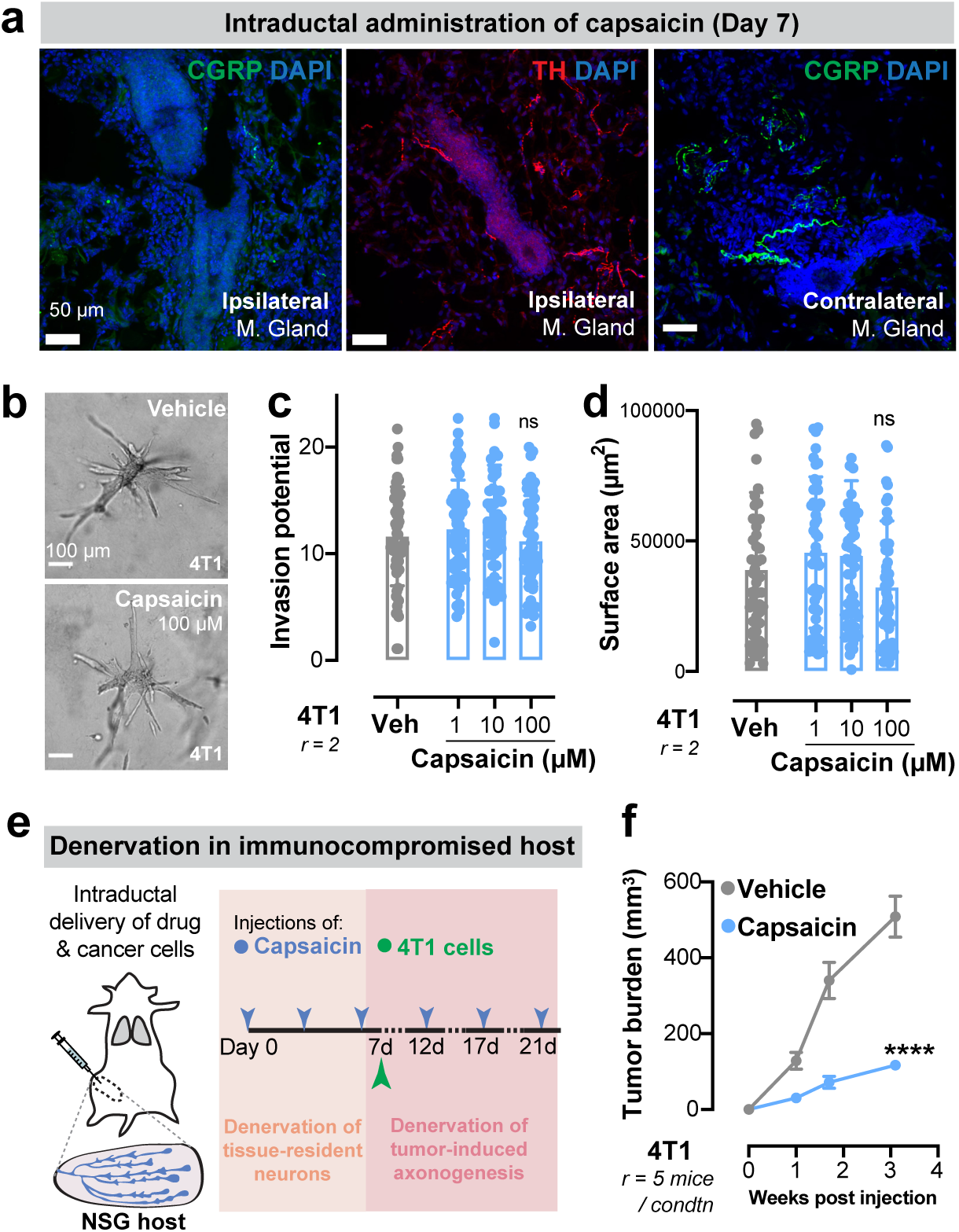
Capsaicin does not alter cancer cell growth or invasion *in vitro* and decreases tumour growth in immune-compromised mice *in vivo*. (a) Representative micrographs of murine mammary glands 7 days after intraductal administration of capsaicin (see Fig. 1n). The ipsilateral mammary gland was devoid of sensory innervation but retained sympathetic innervation (TH+) post-capsaicin treatment. Sensory innervation of the contralateral mammary gland was unaffected. Scale bar, 50 μm. (b) Representative micrographs of 4T1 cancer cell spheroids cultured in the presence of capsaicin (1-100 μM) in the culture medium. Scale bar, 100 μm. (c-d) There is no significant change in the invasiveness (c) or surface area (d) of 4T1 spheroids cultured in the presence of capsaicin. Mean ± SD. ^ns^*p* > 0.9999 (c), *p* = 0.7108 (d) by Kruskal-Wallis test. (e) Schematic of sensory-specific denervation of 4T1 tumours using capsaicin in immune-compromised NSG mice. Capsaicin was injected intraductally to remove tissue-resident innervation prior to injection of 4T1 cancer cells. Capsaicin injections were continued intraductally to sustain the denervation. (f) Capsaicin-treated 4T1 tumours are significantly smaller than vehicle treated tumours when implanted in NSG mice. Mean ± SEM. \*\*\*\**p* < 0.0001 by two-sided t test.

**Extended Data Figure 6:**
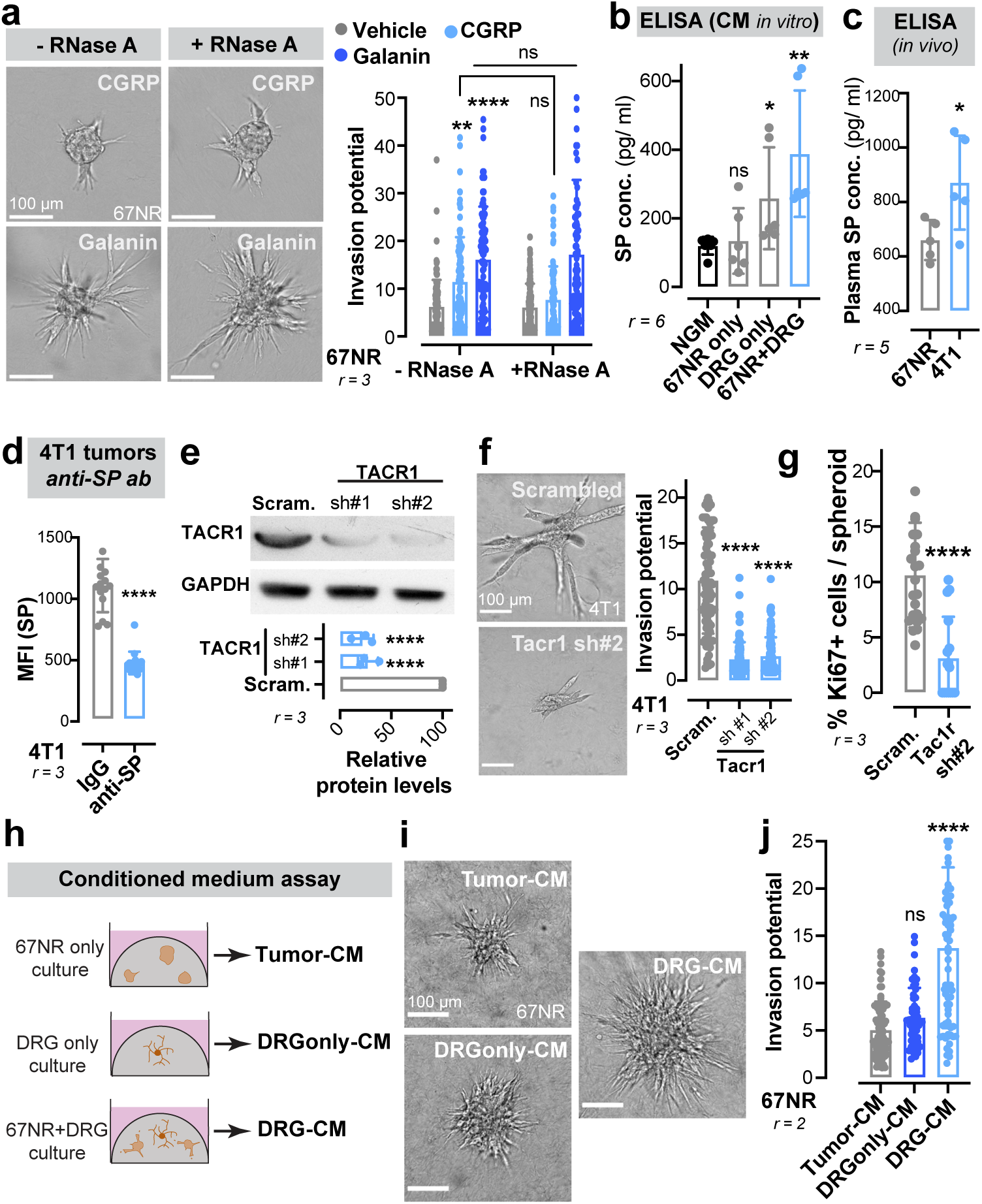
DRG-secreted substance-P drives breast cancer metastasis via the activation of the TACR1 receptor. (a) Left: Representative micrographs of 67NR spheroids cultured in the presence of neuropeptides CGRP or galanin and/ or RNase A. Scale bar, 100 μm. Right: Galanin and CGRP independently promote the invasiveness of 67NR spheroids. RNase A treatment does not significantly affect the ability of either neuropeptide in promoting spheroid invasion. Scale bar, 100 μm. Mean ± SD. \*\**p* = 0.0011, \*\*\*\**p* < 0.0001, ^ns^*p* (CGRP) = 0.1066, ^ns^*p* (galanin) > 0.9999 by Kruskal-Wallis test. (b) There is significantly more substance-P (SP) secreted in the medium of a 67NR-DRG co-culture compared to a culture of either 67NR or DRGs alone. p values are based on individual comparisons with base medium (NGM). Mean ± SD. ^ns^*p* = 0.7281, *p = 0.0476, \*\**p* = 0.0054 by unpaired two-tailed t-test. (c) Plasma SP levels are significantly higher in 4T1 tumour bearing mice relative to mice with 67NR tumours. Mean ± SD. \**p* = 0.0359 by unpaired two-tailed t-test. (d) Mean fluorescence intensity of SP in 4T1 primary tumours treated with an anti-SP antibody is significantly lower compared to those treated with IgG-treated controls. Mean ± SD. \*\*\*\**p* < 0.0001 by Mann-Whitney test. (e) Representative Western blot showing reduced TACR1 levels in 4T1 cancer cells transduced with *Tacr1*-targeting short hairpins (sh#1 and sh#2) relative to a scrambled control. \*\*\*\**p* < 0.0001 by one-way ANOVA. (f-g) Depletion of TACR1 significantly decreases 4T1 spheroid invasiveness (f) and proliferation (g). Scale bar, 100 μm. Mean ± SD. \*\*\*\**p* < 0.0001 by Mann-Whitney test. (h) Conditioned medium was isolated from a 3D collagen I-based culture of 67NR spheroids (tumour-CM), DRG neurons (DRG only-CM), or a 67NR-DRG co-culture (DRG-CM). (i) Representative micrographs of 67NR spheroids cultured in the presence of tumour-CM, DRG only-CM, or DRG-CM. Scale bar, 100 μm. (j) 67NR spheroids cultured with DRG-only CM are not any more invasive compared to those cultured with tumour-CM. In contrast, treatment with DRG-CM significantly increases the invasiveness of 67NR spheroids. Mean ± SD. ^ns^*p* = 0.1463, \*\*\*\**p* < 0.0001 by Kruska-Wallis test.

**Extended Data Figure 7:**
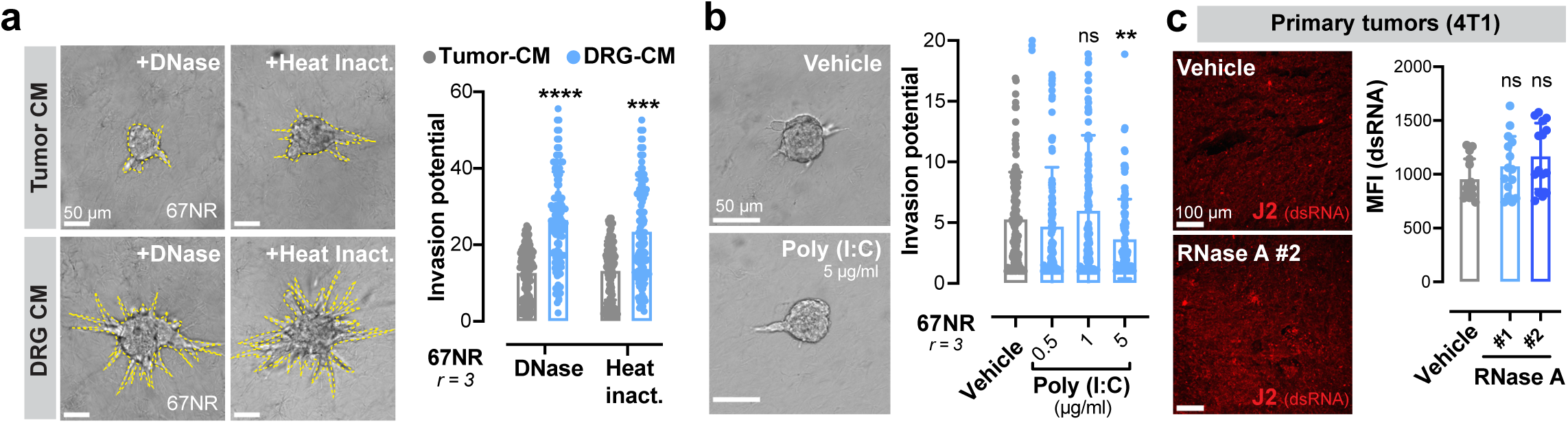
Secreted DNA, dsRNA, or proteins in DRG-CM are not mediators of its pro-metastatic functions. (a) Left: Representative micrographs of 67NR spheroids cultured in the presence of either DNase-treated or heat inactivated tumour-CM or DRG-CM. Scale bar, 50 μm. Right: Neither DNase treatment nor heat inactivation of DRG-CM alters its ability to promote invasiveness of 67NR spheroids. Mean ± SD. \*\*\**p =* 0.001, \*\*\*\**p* < 0.0001 by Kruskal-Wallis test. (b) Treatment with a dsRNA mimetic, Poly (I:C) moderately inhibits the invasion of 67NR spheroids. Scale bar, 50 μm. ^ns^*p* = 0.6948, \*\**p* = 0.0072 by Kruskal-Wallis test. (c) Left: Representative confocal images of 4T1 primary tumours treated with vehicle or RNase A (as described in Fig. 3f) and immunostained for an anti-dsRNA antibody (J2). Right: There is no change in the amount of dsRNA in 4T1 tumours treated with RNase A. Mean ± SD. ^ns^*p* = 0.22 (RNase A #1), 0.0625 (RNase A #2) by one-way ANOVA.

**Extended Data Figure 8:**
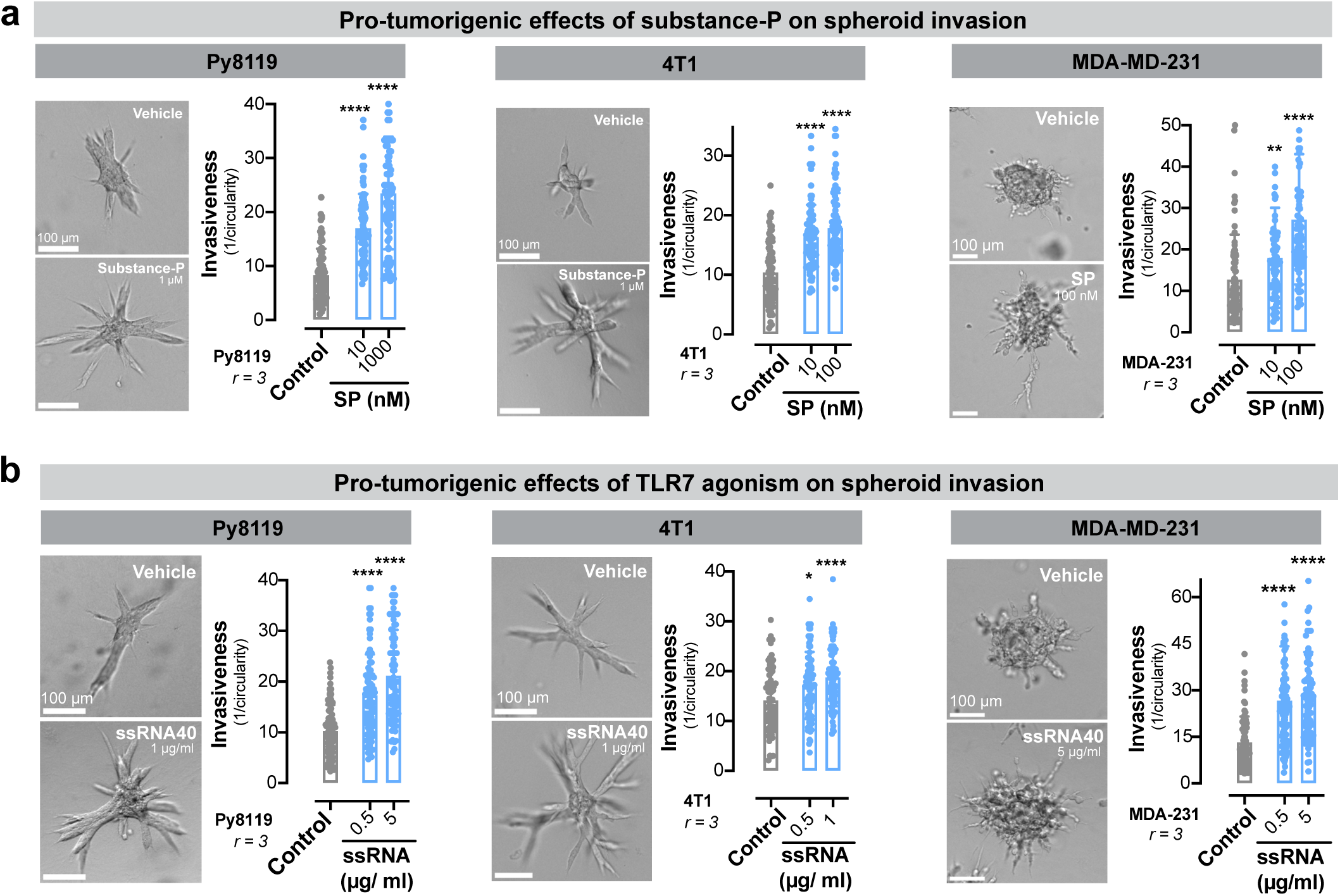
ssRNA-driven activation of TLR7 and SP-driven activation of TACR1 promote breast cancer invasiveness across multiple models of breast cancer. (a) SP significantly increases invasion of Py8119, 4T1, and MDA-MB-231 breast cancer spheroids. Scale bar, 100 μm. Mean ± SD.\*\**p* = 0.0013, \*\*\*\**p* < 0.0001 by Kruskal-Wallis test. (b) A ssRNA mimetic increases invasion of Py8119, 4T1, and MDA-MB-231 breast cancer spheroids. Scale bar, 100 μm. Mean ± SD. \**p* = 0.0235, \*\*\*\**p* < 0.0001 by Kruskal-Wallis test.

**Extended Data Figure 9:**
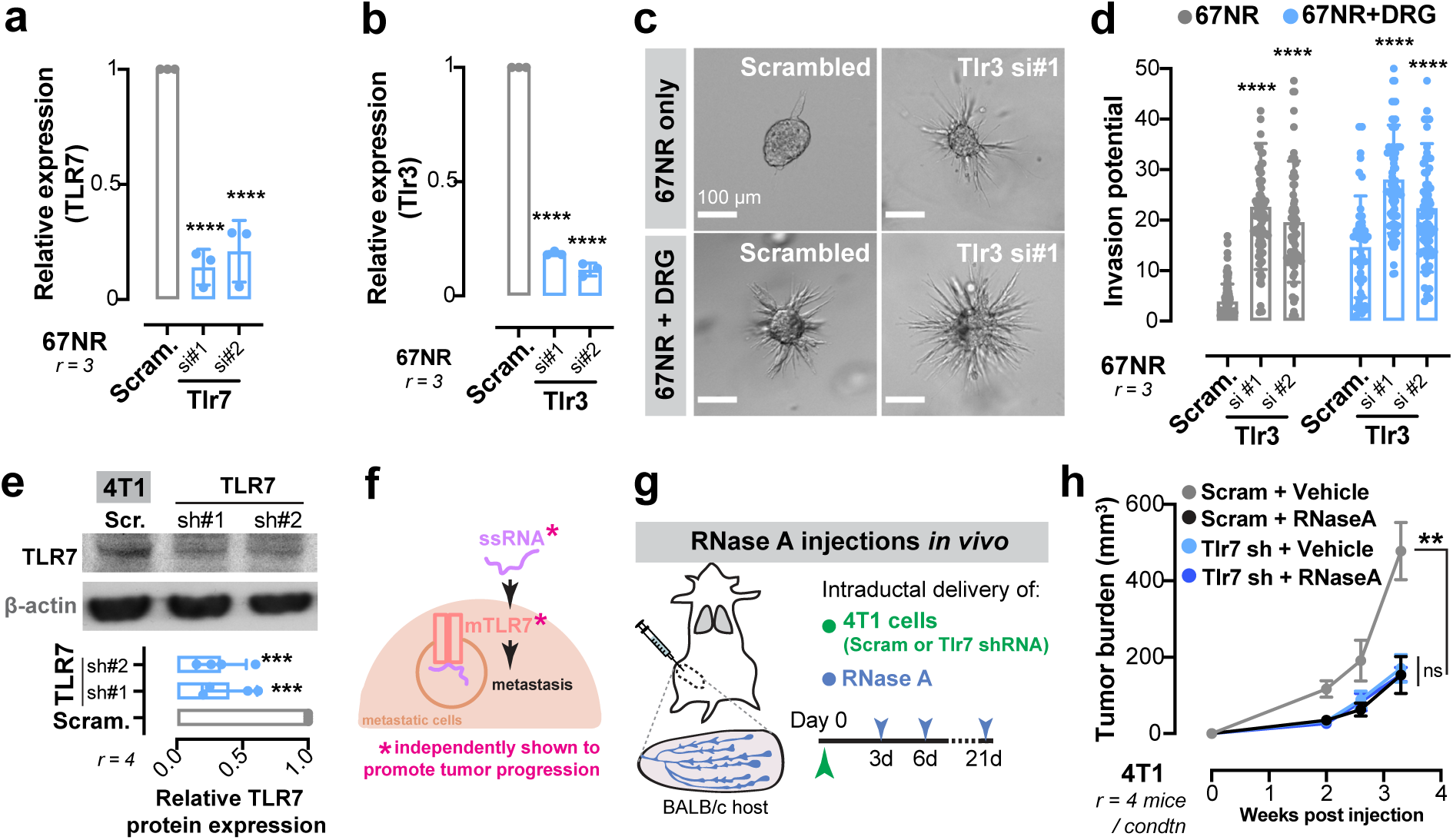
ssRNA-mediated activation of Tlr7 promotes breast cancer invasion and metastasis. (a-b) qPCR validation showing successful depletion of Tlr7 (a) and Tlr3 (b) mRNA in 67NR spheroids after transfection with corresponding targeting siRNAs. Mean ± SD. \*\*\*\**p* < 0.0001 by one-way ANOVA. (c) Representative micrographs of 67NR spheroids transfected with either scrambled or Tlr3 targeting siRNA cultured in the absence (top row) or presence (bottom row) of DRG neurons. Scale bar, 100 μm. (d) Tlr3 depletion significantly increases the invasiveness of 67NR spheroids. Culture of 67NR spheroids knocked down for Tlr3 remain invasive when co-cultured with DRG neurons. Mean ± SD. \*\*\*\**p* < 0.0001 by Kruskal-Wallis test. (e) Representative Western blot depicting lower TLR7 protein levels in 4T1 cancer cells transduced with *Tlr7* shRNA relative to scrambled control. Mean ± SD. \*\*\**p* = 0.0006 by one-way ANOVA. (f) We have independently shown that ssRNAs and TLR7 promote breast cancer metastasis. (g) 4T1 cancer cells transduced with scrambled or Tlr7 shRNA were injected intraductally into BALB/c mice. This was followed by periodic injections with either vehicle or RNase A. (h) RNase A and Tlr7 depletion independently impaired 4T1 tumour growth. However, RNase A treatment of Tlr7 depleted tumours did not further decrease tumour growth, suggesting that ssRNA and Tlr7 function in a common pathway to drive breast cancer tumourigenesis. Mean ± SD. \*\**p* = 0.0033 by one-way ANOVA.

**Extended Data Figure 10:**
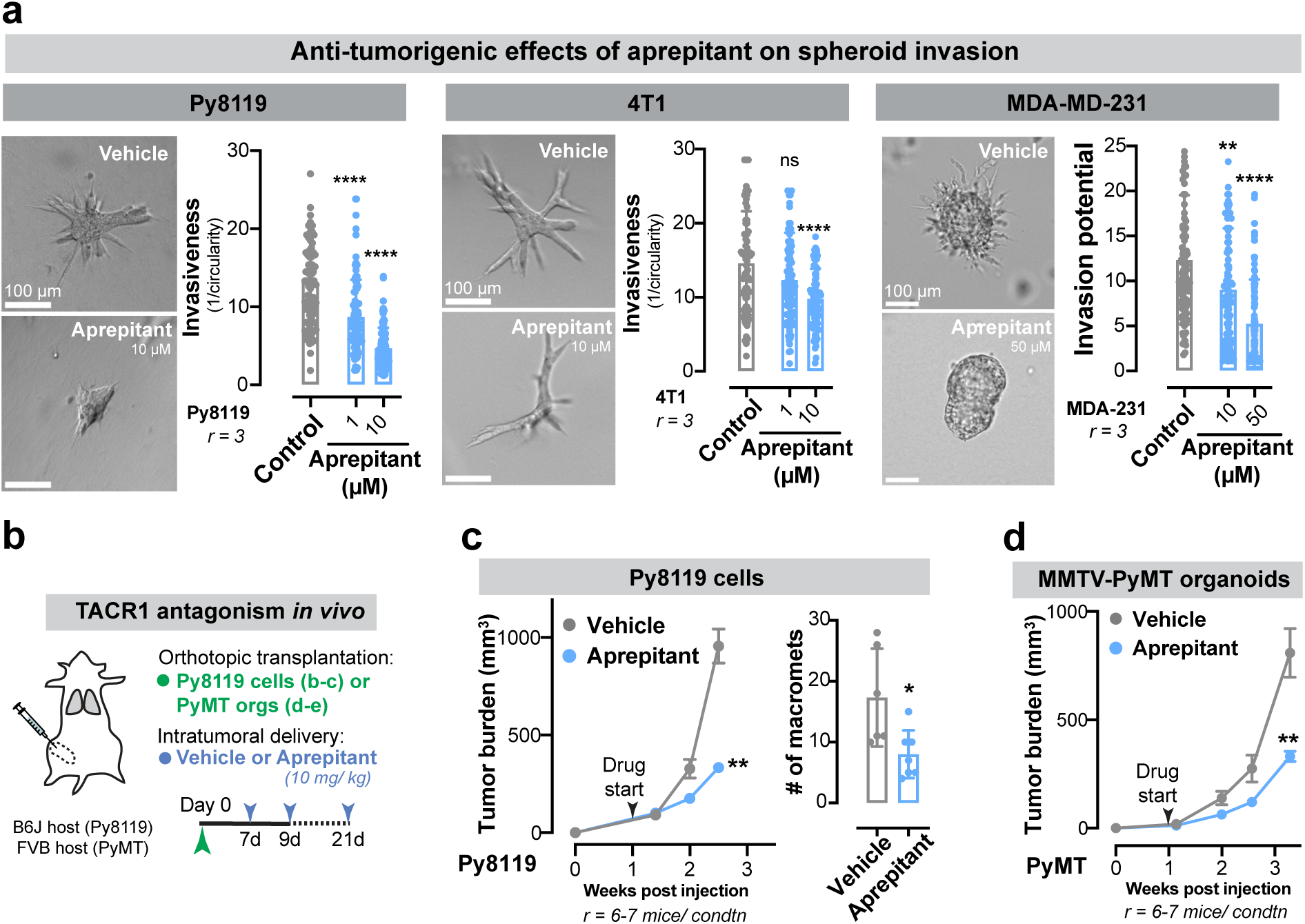
Aprepitant significantly impairs tumour growth and metastasis of Py8119 and MMTV-PyMT models of breast cancer. (a) TACR1 antagonism using aprepitant significantly decreases invasion of Py8119, 4T1, and MDA-MB-231 breast cancer spheroids. Scale bar, 100 μm. ^ns^*p* = 0.9621, \*\**p* = 0.0013, \*\*\*\**p* < 0.0001 by Kruskal-Wallis test. (b) Schematic of TACR1 antagonism using aprepitant in mice with Py8119 and MMTV-PyMT-derived tumours. Cancer cells were injected intraductally into syngeneic host mice followed by periodic intra-tumoral injections of aprepitant. (c) Aprepitant significantly impairs tumour growth and metastastatic colonization of Py8119 cells. (d) Aprepitant significantly impairs growth of MMTV-PyMT tumours.

## MATERIALS AND METHODS

### Animal studies

All animal experiments were conducted in accordance with the Institutional Animal Care and Use Committee at The Rockefeller University. Wild-type BALB/cJ, C57BL/6J, FVB, and *Tac1*-KO (JAX, 004103) animals were obtained from Jackson laboratories. MMTV-PyMT^50^ mice were a generous gift from Kevin J. Cheung. Fixed C3(1)-Tag^51^ tumours were donated by Andrew J. Ewald.

### Primary DRG culture

Dorsal root ganglia (DRG) neurons were isolated from 5–8-week-old female syngeneic mice (BALB/cJ, FVB, or C57BL/6J). Each mouse was perfused with 20 ml of cold HBSS (Sigma-Aldrich, 55021C). Ganglia were collected into cold HBSS, pelleted by spinning at 100 g for 2 minutes. Ganglia were then digested in papain (Worthington LS003126) and collagenase (Worthington LS004176) at 37°C. The digested neurons were triturated with fire-polished Pasteur pipettes. DRG neurons were further purified using a Percoll (GE Healthcare, 17-0891-01) gradient (12.5% and 28%). Pelleted neurons were resuspended in neuronal growth media (DMEM-F12 with HEPES (Fisher Scientific 11-330-032), penicillin-streptomycin (ThermoFischer Scientific 15140-122), bovine serum albumin (BSA; Sigma-Aldrich, A9576), and fetal bovine serum (FBS; Sigma-Aldrich, F4135)) at a concentration of 50-100 neurons/ μl. For all co-culture experiments, the DRG: cancer cell ratio of 1:200 was used unless otherwise specified.

### Isolation of primary tumour organoids

Fresh human tumour samples were received in basal RPMI at 4°C within 24 hours post-surgery and processed as previously described^52^. Mouse mammary tumours were dissected immediately prior to processing. Samples were mechanically disrupted with a scalpel, and enzymatically (collagenase-based) digested. The digested tissue was centrifuged for 10 minutes at 1500 rpm. The pelleted tissue was then treated with DNAse (Sigma, D4263). Differential centrifugation (quick spins at 400 g for 3-7 seconds) was used to separate out single/ stromal cells from epithelial organoids.

### Tissue culture and spheroid generation

4T1, EO771, HEK293T, Py8119 and MDA-MB-231 cells were obtained from American Tissue Type collection (ATCC). 4T07 and 67NR cells were a generous gift from W. P. Schiemann. EO771 cells were maintained in RPMI with 10% FBS and 10 mM HEPES (ThermoFisher Scientific 15-630-080). 67NR, 4T1, and MDA-MB-231 cancer cells were maintained in DMEM (Gibco, 11995065) with 10% FBS. Py8119 cells were cultured in Ham’s F-12K (Kaighn’s) medium (ThermoFisher Scientific, 21127022) with 10% FBS. Spheroids were formed using the hanging drop method^53^. 50, 3000, and 1500 cells from 4T1, 67NR, and Py8119 were resuspended in 25 μl of media and plated into a lid of a tissue-culture treated dish. Droplets were placed sufficiently apart to not merge when lid was inverted and placed back on the PBS-filled reservoir. Spheroids were harvested after an overnight incubation. Spheroids were from 5000 MDA-MB-231 cells per 25 μl of media containing 5% Matrigel (Corning, 354320). Contamination with mycoplasma was ruled out on a quarterly basis using PCR-based protocols^54^.

### Three-dimensional cultures

For invasion assays, organoids/ spheroids were embedded in neutralized, cold-polymerized, fibrillar rat-tail collagen I (Corning, 354236). Invasion assays in the presence of DRG neurons consisted of 100 neurons/ well in a 24-well plate. For colony formation assays using MMTV-PyMT mouse model, organoids were briefly dissociated into small cell clusters and embedded in Matrigel (Corning, 354320; 5,000 clusters/ well in a 24-well plate). For cell lines, colony formation assays were performed using 67NR cancer cells (10,000 cells/ well in a 24-well plate) and DRG neurons (100 neurons/ well in a 24-well plate). The base medium for all organoid cultures (and co-cultures) included DMEM (Gibco, 11995065), L-glutamine (ThermoFisher, 35050061), 1% penicillin-streptomycin (Gibco, 15070063), 30% BSA (Sigma-Aldrich, A9576), FBS (Sigma-Aldrich, F4135), N2 and B27 supplements (ThermoFisher Scientific, 17502048 and 17504001 respectively). Human organoids were additionally supplemented with hEGF ((Sigma-Aldrich, E9644) while murine cultures were supplemented with FGF2 (Sigma, F0291). Conditioned medium was collected 24 and 48 hours after the setup of the original 3D culture in collagen I. Conditioned medium was isolated every 24 hours from 3D cultures in collagen I setup using neuronal growth medium. For conditioned medium experiments, RNase A (ThermoFisher Scientific, EN0531) was used at a concentration of 25 μg/ ml, RNase T1 (ThermoFisher Scientific, EN0541) was used at concentration of 2000 U/ ml, RNase III (ThermoFisher Scientific, AM2290) was used at concentration of 2.5 U/ ml, DNase I was used at a concentration of 2 U/ ml, and heat inactivation was performed at 100°C for 20 minutes.

### Lentiviral production and transduction

Third generation lentivirus system was used to produce virus from HEK293T cells grown in 10 cm plates. HEK293T cells were plated at 70% confluency the night before transfection and cultured in DMEM (Gibco, 11995065) with 10% FBS (Sigma-Aldrich, F4135). Cells were transfected with 2.5 μg of Gag-Pol, 5 μg of VSV-G, 2.5 μg of RSV-Rev and 7.5 μg of pLKO.puro1 vector cloned to contain the appropriate shRNA, using 60 μl of Lipofectamine 2000 (ThermoFisher Scientific, 11668019). After an overnight incubation, the medium was replaced with fresh medium. Virus-containing medium was collected 48- and 72-hours post-transfection. The supernatant was filtered through a 0.45 μm filter (Pall, 4614), mixed 1:1 with fresh medium and 8 μg/ ml polybrene (Sigma-Aldrich, TR-1003-G) to transduce pre-plated 4T1 cells at 50% confluency. Selection with 3 μg of puromycin (ThermoFisher Scientific, A1113803) was conducted 36 hours post-transduction. Knockdown was validated using qPCR and Western Blotting.

Tlr7 shRNA#1 (mouse):

F 5’-CCGGTCTATGGAGAGCCGGTGATAACTCGAGTTATCACCGGCTCTCCATAGATTTTTG-3’ R 5’AATTCAAAAATCTATGGAGAGCCGGTGATAACTCGAGTTATCACCGGCTCTCCATAGA-3’

Tlr7 shRNA#2 (mouse):

F 5’-CCGGACCACTTTGCCACCTAATTTACTCGAGTAAATTAGGTGGCAAAGTGGTTTTTTG-3’ R 5’AATTCAAAAAACCACTTTGCCACCTAATTTACTCGAGTAAATTAGGTGGCAAAGTGGT-3’

Tacr1 shRNA#1 (mouse):

F 5’-CGGGAGGACAGTGACCAATTATTTCTCGAGAAATAATTGGTCACTGTCCTCTTTTTG-3’ R 5’AATTCAAAAAGAGGACAGTGACCAATTATTTCTCGAGAAATAATTGGTCACTGTCCTC-3’

Tacr1 shRNA#2 (mouse):

F 5’-CCGGTGGAGAAGGCAAGCGTTATATCTCGAGATATAACGCTTGCCTTCTCCATTTTTG-3’ R 5’AATTCAAAAATGGAGAAGGCAAGCGTTATATCTCGAGATATAACGCTTGCCTTCTCCA-3’

### siRNA Transfection

Transient knockdown of Tlr7 was performed using at least two independent siRNAs against these genes in 67NR cells (sequences below). The siRNA and Lipofectamine^TM^ RNAiMAX transfection reagent (ThermoFisher Scientific, 13778150) were diluted in Opti-MEM (ThermoFisher Scientific, 31985070) and added to 67NR cells at ∼60% confluency. After an overnight incubation, the culture medium was replaced with fresh medium. Successful knockdown was validated using qPCR and Western Blotting.

Tlr7 (mouse) siRNA sequences:

5’-GCAGAAGGAAUUACUCACAUGCUAA-3’

5’-CCUUUCAGUGAAUAAGAUAUCUCCT-3’

### Protein Isolation

Cell lines were washed with and scraped into ice-cold PBS (without any Ca^2+^ or Mg^2+^), centrifuged for 5 minutes at 300 g (4°C). The cell pellet was resuspended in an appropriate volume of RIPA-based lysis buffer, vortexed and left on ice for 5 to 10 minutes. For primary tumours, ∼10 mg of tumour tissue was homogenized in ∼100 µl ice-cold RIPA lysis buffer using an electric homogenizer (Bel-Art Homogenizer System, Fisher Scientific, 03-421-215; Bel-Art Pestles and Tubes, Fisher Scientific, 03-421-221). Samples were centrifuged at 10,000 g (4°C) for 10 minutes and the supernatants were transferred to new, prechilled Eppendorf tubes. A BCA assay kit (Thermo Scientific, PI23225) was used to quantify the amount of protein in each sample. Samples were stored at -80°C.

### Western blotting

Whole cell protein lysates were thawed on ice for 30 minutes before use. Samples were diluted with 4x protein loading buffer (Li-Cor, 928-40004) and 10x sample reducing agent (ThermoFisher Scientific, NP0009). Equal amounts of protein were loaded in 4-12% Bis-tris Mini Protein Gels (Fisher Scientific, NP0336PK2). SDS-Page was performed in MES-SDS running buffer (20x, ThermoFisher Scientific, NP0002) at 140 V for ∼1 hour or until the dye front had run off the gel. The gels were transferred at 100 V for 1.5 hours at 4°C in transfer buffer (ThermoFisher Scientific, NP0006) onto PVDF membranes (ThermoFisher Scientific, 88520). Membranes were blocked in 5% nonfat dry milk (BioRad, 1706404) in TBS-T (TBS (Cell Signaling Technology, 12498S) + 0.1% Tween (Millipore, P2287)) for 1 hour at room temperature. Primary antibodies were diluted in 3% BSA/ TBS-T (Millipore, A2153) and incubated overnight at 4°C. Membranes were washed thrice with 0.1% TBS-T and incubated for 1 hour at room temperature with corresponding HRP-linked secondary antibodies, added at a dilution of 1:10,000. Membranes were then washed three times with 0.1% TBS-T and activated by a ECL blotting solution (ThermoFisher Scientific, 32106). X-Ray films (Imaging Solutions Company, 110102) were exposed to the membrane 10 seconds up to 2 minutes and developed afterwards. ImageJ was used to quantify band intensity. Primary antibodies used included mouse βIII-tubulin (1:1000; Biolegend, MMS-435P, Clone TUJ1), rabbit GAPDH (1:1000; Cell Signaling 2118, 14C10), mouse HSP-60 (1:1000; Santa Cruz Biotechnology, sc-13115), and rabbit TACR1 (1:1000; ThermoFisher, PA1-32229).

### Immunofluorescence

Tumours, lungs, and mammary glands harvested from mice were fixed in 4% paraformaldehyde (Fisher Scientific, AA433689M) overnight at 4°C and then transferred into 25% sucrose (Sigma-Aldrich, S0389), 0.1% sodium azide (Fisher Scientific, 71448-16) in PBS for 24 hours at 4°C. Tissues were embedded in Tissue-Tek O.C.T. Compound (VWR, 4583) and frozen at -80°C. Sections (20-50 µm thick) were cut using a cryostat set to -20°C cutting temperature for lungs and tumours and -27°C for mammary glands. The sections were collected onto Superfrost Plus microscope Slides (Fisher Scientific, 12-550-15). Prior to immunofluorescent staining, O.C.T. was removed by incubating the slides in PBS for 1 hour.

Formalin-fixed paraffin-embedded tissue sections were dewaxed and rehydrated using xylene and descending concentrations of ethanol. Antigen retrieval was performed by boiling samples in citrate buffer (Sigma-Aldrich, C9999) for 30 minutes. Sections were then permeabilized using 0.5% Triton-X-100 (Millipore, T9284). Slides were blocked in 10%FBS (Sigma-Aldrich, F4135), 1%BSA (Millipore, A2153), 0.1% Tween-20 (Millipore, P2287) in PBS for up to 2 hours at room temperature. Incubation with primary and secondary antibodies (diluted in 1% FBS, 1% BSA, 0.1% Tween-20) was performed overnight at 4°C or for 2-3 hours at room temperature. Slides were mounted with Prolong Gold (Fisher Scientific, P36930). Primary antibodies used in the study include rabbit anti-CGRP (1:250, Cell Signaling, mAb#14959), mouse anti-βIII-tubulin (1:200, Biolegend, MMS-435P, Clone TUJ1), rabbit anti-Ki-67 (abcam, ab15580), rabbit anti-substance P (Sigma-Aldrich, AB1566), mouse anti-neurofilament-L (Cell Signaling, #2837), mouse anti-dsRNA (1:200, Axxora, JBS-RNT-SCI-10010200) and phalloidin (1:200, ThermoFisher Scientific, A12379). Nuclei were counterstained using Hoechst (1:500, ThermoFisher Scientific, H3579). All secondary antibodies were AlexaFluor conjugates (1:200, ThermoFisher Scientific).

### qPCR

RNA was extracted from cells using the RNA purification kit (Norgen Biotek, 37500)), according to the manufacturer’s protocol. Reverse transcription (RT-PCR) was performed on 2-5 μg of RNA using the SuperScript III First-Strand Synthesis System (ThermoFisher Scientific, 18080051). Real time qPCR was conducted in a 384-well PCR microplate (Fisher scientific, AB-1384) performed on an Applied Biosystem StepOne Real-Time PCR System using the Fast SYBR Green master mix (Fisher Scientific, 43-856-18). ΔCt values were used to calculate the relative levels of the mRNA of interest which were normalized to the mRNA levels of GAPDH. Primers used in the studies include:

GAPDH (mouse): Forward – 5’-AGGTCGGTGTGAACGGATTTG-3’ Reverse – 5’-TGTAGACCATGTAGTTGAGGTCA-3’

Tlr7 (mouse): Forward – 5’-CACCACCAATCTTACCCTTACC-3’ Reverse – 5’-CAGATGGTTCAGCCTACGGAA-3’

Tacr1 (mouse): Forward – 5’-CTCCACCAACACTTCTGAGTC-3’ Reverse – 5’-TCACCACTGTATTGAATGCAGC-3’

### Mammary-fat-pad injections

Cancer cells (67NR, 4T1, EO771, or Py8119) were resuspended in a 50:50 mix of PBS: Matrigel (Corning, 354320) at a concentration of 10^6 cells/ ml. For DRG co-transplantations, 100 DRGs were injected per animal (added to the cancer cell suspension just prior to injections into the animal). Orthotopic transplantations were conducted in 5-7-week-old syngeneic female mice as previously described^55^. Mice were anesthetized, immobilized, and the surgical site was shaved and sterilized using povidone iodine (Abcam, ab143439). A midline incision was made to expose the abdominal mammary gland. 50 μl of the cell suspension was injected into the mammary gland. The surgical wound was closed using 9 mm autoclips. Tumours measurements were taken using digital calipers and the total tumour burden was not allowed to exceed 1500 mm^3^. Tumours and lungs were typically harvested 3-4 weeks after transplantation. For CTC enumeration, cardiac puncture was used to obtain 750 μl – 1 ml of blood from animals with late-stage tumours. Following lysis of red blood cells, the remainder of the sample was smeared on a glass microscope slide and examined under a confocal microscope for mCherry+ expression. Cells with intact nuclei and mCherry expression were identified as a CTC.

### Tail-vein injections

67NR cells were resuspended in PBS at a concentration of 750,000 cells/ ml and stored on ice. Tail vein injections were performed in 6-9-week-old syngeneic female mice. Mice were placed in a heating chamber set to 37°C for 10 minutes and immobilized in a restraining device. 200 μl of the cell suspension was injected via the tail vein. Lungs from these mice were harvested ∼1 week later and the number of macro-metastases were examined under the dissection microscope

### Intraductal injections

Intraductal injections of cancer cells or reagents/ drugs were performed to restrict delivery within the ductal epithelium. Injections were performed under a dissection microscope as previously described^56^. Using a pair of micro-dissecting tweezers, the dead skin around the nipple is removed. A 33-gauge micro-syringe (needle: CAL7637-01, syringe: 89221-012) is used to inject no more than 20 μl into the ductal epithelium. 4T1 cancer cells were resuspended at a concentration of 10^6 cells/ ml in PBS. Capsaicin was dissolved in 10% ethanol, 10% Tween-80, and 80% saline and injected into mice (20 μl/ mouse) at a concentration of 75 mg/ kg. Anti-SP (BioGeneX AR069GP) or IgG control (BioXCell BE0095) antibodies were injected into 4T1 tumour-bearing NOD-SCID gamma mice at a concentration of 20 μg/ mouse. RNase A (two sources: ThermoFisher Scientific EN0531 and 12091021) was used at a concentration of 5 mg/ ml.

### Genotyping of transgenic mouse lines

Tail snips collected from 2.5-week-old mice were lysed in DirectPCR Lysis Reagent (Viagen Biotech, 102-T) and Proteinase K (Millipore Sigma, 3115828001) to extract DNA. PCR reactions using the primers listed below were used to detect the presence of a transgene.

MMTV-PyMT:

F: 5’-GGAAGCAAGTACTTCACAAGGG-3’ R: 5’-GGAAAGTCACTAGGAGCAGGG-3’

Internal positive control F: 5’-CAAATGTTGCTTGTCTGGTG-3’ Internal positive control R: 5’-GTCAGTCGAGTGCACAGTTT-3’ Positive control runs at 200 bp while the transgene runs at 556 bp.

### Tac1-KO

F: 5’-AGAATTTAAAGCTCTTTTGCC-3’

R: 5’-GCTCATCAGTATGTGACATAGAAA-3’

Mutant allele runs at 175 bp while the wild-type runs at 190 bp.

### Optical tissue clearing

Clearing of murine mammary tumours was performed using a previously described AdipoCLEAR protocol^57^. Tumours were fixed in 4% paraformaldehyde (ThermoFisher Scientific, AA433689M) and washed in increasing concentrations of methanol (20-100%) in B1N buffer (water, 0.1% Triton X-100 (Sigma-Aldrich, T9284), 0.3 M glycine (Sigma-Aldrich, G5417)). Delipidation was performed using 100% dicholoromethane (Sigma-Aldrich, 270997). Samples were washed in decreasing concentrations of methanol (100-20%) in B1N buffer. Tumours were incubated with primary antibodies (anti-CRP, 1:200, Cell Signaling, mAb#14959) for 4 days with continuous shaking, washed, and incubated with secondary antibodies (AlexaFluor conjugates) for another 4 days. After immunolabelling, samples were dehydrated in an increasing series of methanol/ water (25-100%). Samples were washed in dicholoromethane overnight and cleared in dibenzyl ether (Sigma-Aldrich, 33630).

### Imaging and image analysis

Optically cleared tissues were imaged on a light sheet microscope (LaVision BioTec) with a 4x/0.3 LVMI-Fluar objective with 5.6-6 mm WD. Images were imported into Imaris and rendered in 3D. Confocal microscopy was performed using either the LSM 780 laser scanning confocal microscope (Zeiss) or A1R confocal microscope (Nikon). Phase contrast/ brightfield images using the CellDiscoverer7 (Zeiss). For experiments in which immunofluorescence was used to infer relative protein expression levels, all images were collected and processed using identical parameters. All image analysis was performed using Fiji/ ImageJ2. Extent of innervation in tissue sections was estimated by manually counting the number of nerve bundles, positively identified by βIII-tubulin or neurofilament-L. Invasiveness of organoids/ spheroids was calculated as the inverse of the circularity of the structure. For substance-P expression levels in human breast tumours, fluorescence intensity was measured within an ROI drawn around any cancer-cell rich areas of the tumour.

### Statistical analysis

For all experiments, independent biological replicates are listed as “r”. Unless otherwise noted, all data are expressed as mean ± standard deviation. Groups were compared using tests for significance as indicated in the figure legends and the text. A significant difference was concluded at *P* < 0.05.

### RNA-sequencing

Libraries were constructed from 500 ng of total RNAs isolated from 4T1 (scrambled vs Tlr7 shRNA) using the TruSeq RNA Library Prep Kit (Illumina). Constructed libraries were sequenced using Illumina NextSeq (High Output, 75 SR) at the Rockefeller University Genomics Center. Sequencing reads were mapped to the mouse genome (assembly GRCm38) using STAR aligner (v2.7.5) at default settings. STAR was also used for counting reads mapping to genes. Further analyses were performed using R (v4.1.0). Differentially expressed genes were determined using DESeq2 (v1.32.0). For gene set enrichment analysis, genes were ranked according to log fold changes shrunken with the ‘ashr’ method and enriched pathways of the Kegg database were identified using the ‘gseKEGG’ function of the clusterProfiler package for R (v4.0.0).

### Analysis of the TCGA and METABRIC studies

To assess the association of TLR7-dependent signaling with outcome in breast cancer patients, we analyzed the association of a TLR7 (orthologous to human TLR8)-dependent gene signature with survival in two large independent datasets. Specifically, we generated a signature of genes downregulated in cancer spheroids in which Tlr7 expression was silenced (see section above) and calculated the TLR7 signature score by summing the scaled expression of the genes contained in the signature and using the ranked signature score as a covariate in a Cox proportional hazards model including tumour stage, age and the ranked signature score using the Survival (v3.2-11) and forestmodel (v0.6.2) packages for R. For the TCGA study^1^, we used clinical data as recently curated^2^ and included only primary tumour samples. For the METABRIC study we used the validation cohort^3^.

### Data availability

Raw sequencing data and count tables for newly generated sequencing data will be made available via public repositories before publication. Data for the METABRIC and TCGA studies are publicly available under the following accession numbers: METABRIC, EGA accession EGAS00000000083; TCGA, dbGaP accession phs000178.v10.p8.

### Code availability

All computer code is available from the corresponding author upon request.

## REFERENCES

1 Ayala, G. E. et al. Cancer-related axonogenesis and neurogenesis in prostate cancer. Clin Cancer Res 14, 7593–7603 (2008). 10.1158/1078-0432.CCR-08-1164

2 Albo, D. et al. Neurogenesis in colorectal cancer is a marker of aggressive tumor behavior and poor outcomes. Cancer 117, 4834–4845 (2011). 10.1002/cncr.26117

3 Huang, D. et al. Nerve fibers in breast cancer tissues indicate aggressive tumor progression. Medicine (Baltimore) 93, e172 (2014). 10.1097/MD.0000000000000172

4 Shao, J. X. et al. Autonomic nervous infiltration positively correlates with pathological risk grading and poor prognosis in patients with lung adenocarcinoma. Thorac Cancer 7, 588–598 (2016). 10.1111/1759-7714.12374

5 Oertel, H. Innervation and Tumour Growth: A Preliminary Report. Can Med Assoc J 18, 135–139 (1928).

6 Shapiro, D. M. & Warren, S. Cancer innervation. Cancer Res 9, 707–711, illust (1949).

7 Young, H. H. On the Presence of Nerves in Tumors and of Other Structures in Them as Revealed by a Modification of Ehrlich’s Method of “Vital Staining” with Methylene Blue. J Exp Med 2, 1–12 (1897). 10.1084/jem.2.1.1

8 Liebig, C., Ayala, G., Wilks, J. A., Berger, D. H. & Albo, D. Perineural invasion in cancer: a review of the literature. Cancer 115, 3379–3391 (2009). 10.1002/cncr.24396

9 Jiang, N. et al. Incorporation of perineural invasion of gastric carcinoma into the 7th edition tumor-node-metastasis staging system. Tumour Biol 35, 9429–9436 (2014). 10.1007/s13277-014-2258-5

10 Zhao, C. M. et al. Denervation suppresses gastric tumorigenesis. Sci Transl Med 6, 250ra115 (2014). 10.1126/scitranslmed.3009569

11 Liebig, C. et al. Perineural invasion is an independent predictor of outcome in colorectal cancer. J Clin Oncol 27, 5131–5137 (2009). 10.1200/JCO.2009.22.4949

12 DeLancey, J. O. et al. Evidence of perineural invasion on prostate biopsy specimen and survival after radical prostatectomy. Urology 81, 354–357 (2013). 10.1016/j.urology.2012.09.034

13 Magnon, C. et al. Autonomic nerve development contributes to prostate cancer progression. Science 341, 1236361 (2013). 10.1126/science.1236361

14 Balood, M. et al. Nociceptor neurons affect cancer immunosurveillance. Nature 611, 405–412 (2022). 10.1038/s41586-022-05374-w

15 Kamiya, A. et al. Genetic manipulation of autonomic nerve fiber innervation and activity and its effect on breast cancer progression. Nat Neurosci 22, 1289–1305 (2019). 10.1038/s41593-019-0430-3

16 Jiang, Y. Q. & Oblinger, M. M. Differential regulation of beta III and other tubulin genes during peripheral and central neuron development. J Cell Sci 103 ( Pt 3), 643–651 (1992). 10.1242/jcs.103.3.643

17 Doran, J. F., Jackson, P., Kynoch, P. A. M. & Thompson, R. J. Isolation of PGP 9.5, a New Human Neurone-Specific Protein Detected by High-Resolution Two-Dimensional Electrophoresis. Journal of Neurochemistry 40, 1542–1547 (1983). 10.1111/j.1471-4159.1983.tb08124.x

18 Györffy, B. et al. An online survival analysis tool to rapidly assess the effect of 22,277 genes on breast cancer prognosis using microarray data of 1,809 patients. Breast Cancer Research and Treatment 123, 725–731 (2010). 10.1007/s10549-009-0674-9

19 Luppi, P. H., Fort, P. & Jouvet, M. Iontophoretic application of unconjugated cholera toxin B subunit (CTb) combined with immunohistochemistry of neurochemical substances: a method for transmitter identification of retrogradely labeled neurons. Brain Res 534, 209–224 (1990). 10.1016/0006-8993(90)90131-t

20 Russell, F. A., King, R., Smillie, S. J., Kodji, X. & Brain, S. D. Calcitonin gene-related peptide: physiology and pathophysiology. Physiol Rev 94, 1099–1142 (2014). 10.1152/physrev.00034.2013

21 Nguyen-Ngoc, K. V. et al. ECM microenvironment regulates collective migration and local dissemination in normal and malignant mammary epithelium. Proc Natl Acad Sci U S A 109, E2595–2604 (2012). 10.1073/pnas.1212834109

22 Aslakson, C. J. & Miller, F. R. Selective events in the metastatic process defined by analysis of the sequential dissemination of subpopulations of a mouse mammary tumor. Cancer Res 52, 1399–1405 (1992).

23 Jancso, G., Kiraly, E., Such, G., Joo, F. & Nagy, A. Neurotoxic effect of capsaicin in mammals. Acta Physiol Hung 69, 295–313 (1987).

24 Bujak, J. K., Kosmala, D., Szopa, I. M., Majchrzak, K. & Bednarczyk, P. Inflammation, Cancer and Immunity-Implication of TRPV1 Channel. Front Oncol 9, 1087 (2019). 10.3389/fonc.2019.01087

25 Rosenfeld, M. G. et al. Production of a novel neuropeptide encoded by the calcitonin gene via tissue-specific RNA processing. Nature 304, 129–135 (1983). 10.1038/304129a0

26 US, V. E. & Gaddum, J. H. An unidentified depressor substance in certain tissue extracts. J Physiol 72, 74–87 (1931). 10.1113/jphysiol.1931.sp002763

27 Tatemoto, K., Rokaeus, A., Jornvall, H., McDonald, T. J. & Mutt, V. Galanin - a novel biologically active peptide from porcine intestine. FEBS Lett 164, 124–128 (1983). 10.1016/0014-5793(83)80033-7

28 Cao, Y. Q. et al. Primary afferent tachykinins are required to experience moderate to intense pain. Nature 392, 390–394 (1998). 10.1038/32897

29 Kastin, A. Handbook of biologically active peptides. (Academic press, 2013).

30 Nishikawa, S. et al. Two histidine residues are essential for ribonuclease T1 activity as is the case for ribonuclease A. Biochemistry 26, 8620–8624 (1987). 10.1021/bi00400a019

31 Robertson, H. D., Webster, R. E. & Zinder, N. D. Purification and properties of ribonuclease III from Escherichia coli. J Biol Chem 243, 82–91 (1968).

32 Heil, F. et al. Species-specific recognition of single-stranded RNA via toll-like receptor 7 and 8. Science 303, 1526–1529 (2004). 10.1126/science.1093620

33 Alexopoulou, L., Holt, A. C., Medzhitov, R. & Flavell, R. A. Recognition of double-stranded RNA and activation of NF-kappaB by Toll-like receptor 3. Nature 413, 732–738 (2001). 10.1038/35099560

34 Lund, J. M. et al. Recognition of single-stranded RNA viruses by Toll-like receptor 7. Proc Natl Acad Sci U S A 101, 5598–5603 (2004). 10.1073/pnas.0400937101

35 Kawasaki, T. & Kawai, T. Toll-like receptor signaling pathways. Front Immunol 5, 461 (2014). 10.3389/fimmu.2014.00461

36 Curtis, C. et al. The genomic and transcriptomic architecture of 2,000 breast tumours reveals novel subgroups. Nature 486, 346–352 (2012). 10.1038/nature10983

37 Hale, J. J. et al. Structural optimization affording 2-(R)-(1-(R)-3, 5-bis(trifluoromethyl)phenylethoxy)-3-(S)-(4-fluoro)phenyl-4- (3-oxo-1,2,4-triazol-5-yl)methylmorpholine, a potent, orally active, long-acting morpholine acetal human NK-1 receptor antagonist. J Med Chem 41, 4607–4614 (1998). 10.1021/jm980299k

38 Hesketh, P. J. et al. The oral neurokinin-1 antagonist aprepitant for the prevention of chemotherapy-induced nausea and vomiting: a multinational, randomized, double-blind, placebo-controlled trial in patients receiving high-dose cisplatin--the Aprepitant Protocol 052 Study Group. J Clin Oncol 21, 4112–4119 (2003). 10.1200/JCO.2003.01.095

39 Renz, B. W. et al. beta2 Adrenergic-Neurotrophin Feedforward Loop Promotes Pancreatic Cancer. Cancer Cell 34, 863–867 (2018). 10.1016/j.ccell.2018.10.010

40 Zahalka, A. H. et al. Adrenergic nerves activate an angio-metabolic switch in prostate cancer. Science 358, 321–326 (2017). 10.1126/science.aah5072

41 Partecke, L. I. et al. Subdiaphragmatic vagotomy promotes tumor growth and reduces survival via TNFalpha in a murine pancreatic cancer model. Oncotarget 8, 22501–22512 (2017). 10.18632/oncotarget.15019

42 Renz, B. W. et al. Cholinergic Signaling via Muscarinic Receptors Directly and Indirectly Suppresses Pancreatic Tumorigenesis and Cancer Stemness. Cancer Discov 8, 1458–1473 (2018). 10.1158/2159-8290.CD-18-0046

43 Gerendai, I. et al. Transneuronal labelling of nerve cells in the CNS of female rat from the mammary gland by viral tracing technique. Neuroscience 108, 103–118 (2001). 10.1016/s0306-4522(01)00399-2

44 Hebb, C. & Linzell, J. L. Innervation of the mammary gland. A histochemical study in the rabbit. Histochem J 2, 491–505 (1970). 10.1007/BF01003127

45 Amit, M. et al. Loss of p53 drives neuron reprogramming in head and neck cancer. Nature 578, 449–454 (2020). 10.1038/s41586-020-1996-3

46 Kalinichenko, V. V., Mokyr, M. B., Graf, L. H., Jr., Cohen, R. L. & Chambers, D. A. Norepinephrine-mediated inhibition of antitumor cytotoxic T lymphocyte generation involves a beta-adrenergic receptor mechanism and decreased TNF-alpha gene expression. J Immunol 163, 2492–2499 (1999).

47 Mohammadpour, H. et al. beta2 adrenergic receptor-mediated signaling regulates the immunosuppressive potential of myeloid-derived suppressor cells. J Clin Invest 129, 5537–5552 (2019). 10.1172/JCI129502

48 Tavora, B. et al. Tumoural activation of TLR3-SLIT2 axis in endothelium drives metastasis. Nature 586, 299–304 (2020). 10.1038/s41586-020-2774-y

49 Benci, J. L. et al. Opposing Functions of Interferon Coordinate Adaptive and Innate Immune Responses to Cancer Immune Checkpoint Blockade. Cell 178, 933–948 e914 (2019). 10.1016/j.cell.2019.07.019

50 Guy, C. T., Cardiff, R. D. & Muller, W. J. Induction of mammary tumors by expression of polyomavirus middle T oncogene: a transgenic mouse model for metastatic disease. Mol Cell Biol 12, 954–961 (1992). 10.1128/mcb.12.3.954-961.1992

51 Maroulakou, I. G., Anver, M., Garrett, L. & Green, J. E. Prostate and mammary adenocarcinoma in transgenic mice carrying a rat C3(1) simian virus 40 large tumor antigen fusion gene. Proc Natl Acad Sci U S A 91, 11236–11240 (1994). 10.1073/pnas.91.23.11236

52 Padmanaban, V. et al. Organotypic culture assays for murine and human primary and metastatic-site tumors. Nat Protoc 15, 2413–2442 (2020). 10.1038/s41596-020-0335-3

53 Foty, R. A simple hanging drop cell culture protocol for generation of 3D spheroids. J Vis Exp (2011). 10.3791/2720

54 Young, L., Sung, J., Stacey, G. & Masters, J. R. Detection of Mycoplasma in cell cultures. Nat Protoc 5, 929–934 (2010). 10.1038/nprot.2010.43

55 Padmanaban, V. et al. E-cadherin is required for metastasis in multiple models of breast cancer. Nature 573, 439–444 (2019). 10.1038/s41586-019-1526-3

56 Krause, S., Brock, A. & Ingber, D. E. Intraductal injection for localized drug delivery to the mouse mammary gland. J Vis Exp (2013). 10.3791/50692

57 Chi, J. et al. Three-Dimensional Adipose Tissue Imaging Reveals Regional Variation in Beige Fat Biogenesis and PRDM16-Dependent Sympathetic Neurite Density. Cell Metab 27, 226–236 e223 (2018). 10.1016/j.cmet.2017.12.011

